# RAD-seq unravels local adaptation along a latitudinal gradient for the wild rice *Zizania latifolia*

**DOI:** 10.1101/2023.11.25.568668

**Authors:** Godfrey Kinyori Wagutu, Xiangrong Fan, Miriam Chepkwemoi Tengwer, Henry Kariuki Njeri, Wei Li, Yuanyuan Chen

## Abstract

The rapid climate change poses considerable threat to biodiversity by constantly altering species distribution, phenotypic variation and allele frequencies. Understanding the interplay between the climate and species phenotypic and genetic variation is thus crucial to inform conservation strategies. In this study, we investigated local adaptation of a widespread aquatic species in China, *Zizania latifolia*.

We performed restriction-site associated DNA sequencing (RAD-seq) for 66 wild samples composed of 10 populations, two populations in each of the five major eco-geographical regions of China and 6 cultivated samples derived from three cultivars found in central China. We assessed genetic diversity and structure, genetic-environmental association, and genome-wide association studies for the 60 wild samples.

Low levels of genetic variability were found in the *Z. latifolia* populations ranging from *H*_E_ = 0.08 to *H*_E_ = 1.52. Population structure showed that samples belonged to two major groups, north and south clades, split along a temperature boundary. It was estimated that the two groups diverged during the 8.2kiloyear event and later experienced severe genetic bottlenecks with advancement of agriculture and increase in human population some 2k years later.

Landscape analysis using SNP and trait data showed that environment was more important in shaping the genetic structure than geography, but the combined effect of environment and geography explained >90% of both genetic and morphological variation. Several loci were found to be correlated with environmental variables as well as morphological traits, most of which were annotated as retrotransposons. Considering the abundance of transposable elements in the *Z. latifolia* genome, differentiation and local adaptation was inferred to be partly driven by temperature-induced transposable elements activity.

## INTRODUCTION

The rapid and continuing climate change poses considerable threats to biodiversity, evidenced by measurable changes in species distribution, phenotypic variation and allele frequencies (Hancock et al., 2011). These alterations are brought about by the need for species to escape demographic collapses by migrating to more suitable locations or adjusting to the changing environment through mutations and innovations from standing genetic variability and phenotypic plasticity (Sang et al., 2022). Given that wetlands are generally isolated and surrounded by expansive terrestrial landscape, migration among different water bodies may be difficult for aquatic species. Alternatively, adapting to the environmental changes becomes a more beneficial means, although sometimes adaptation is difficult to keep up with the changing environment (Xu et al., 2023). Widespread species are generally considered to be more adaptable (Xu et al., 2023). In this case, the question is raised: whether the widespread species can successfully cope with the rapid environmental changes, that is, whether the fate of the widespread species is more optimistic in the future ever-changing environments.

*Zizania latifolia* (Griseb.) Turcz. Ex Stapf, commonly known as Chinese wild rice, is a perennial emergent aquatic plant, widely occurring along lake and rivers margins, ponds and marshes in Eastern Asia (Xu & Zhou, 2017). The plant is a member of Oryzeae rice tribe, and it shares the genus *Zizania* with other three species (*Z. aquatic*, *Z. palustris* and *Z. texana*) from North America. It is anemophilous and reproduces sexually by seeds or asexually by rhizomes. In China, the wild rice has been traditionally used for its seeds and currently for its stem that become swollen and soft after infection with *Ustilago esculenta* fungus (Guo et al. 2007). Additionally, because of the close relationship with rice, the plant has been used to improve rice varieties owing to its superior traits such as flood tolerance and strong stem. It is also used in bioremediation due to its high nutrient and metal uptake capacity (Liu et al., 2007; Peng et al., 2013). Natural *Z. latifolia* populations are distributed in the East China along a wide stretch of latitudinal zones (21^◦^-50^◦^N), which spans five major eco-geographic regions. Each eco-geographic region has different abiotic and biotic factors which could influence species local adaptation and migration (Wu et al., 2003). Therefore, *Z. latifolia* provides a perfect model to study local adaptation through assessment of phenotypic variation and allele frequency across the varying environmental gradient.

With the advancement in sequencing technology, genome scans are increasingly used in studies of adaptive evolution in plants (Lotterhos & Whitlock, 2015). However, some of such studies tend to link genotypes and environment, neglecting the phenotypic traits of plants. This shortcoming can be overcome by genome-wide association study (GWAS, reviewed in Korte & Farlow 2013), where traits measured at the common garden are correlated with allele frequency to identify loci enriched for specific traits. Additionally, most previous studies focused on domesticated species while ignoring natural populations, especially wild-relatives of cultivated crops. With the aggravation of wetland habitat destruction and environmental changes, the molecular basis of adaptation of these natural populations should be understood, which could provide deep insight into local adaptation and inform conservation strategies.

In this study, genetic variation and population divergence were measured using single nucleotide polymorphisms (SNPs) derived from the restriction site-associated DNA sequencing (RAD-seq) of 10 *Z. latifolia* populations from different latitudes and six popular cultivars domesticated in the mid-lower reaches of Yangtze River. We propose two hypotheses as follows: (1) Environmental heterogeneity at different latitudes may exert strong selection pressure on *Z. latifolia*, which could result in obvious population divergence and genetic distinction among populations; (2) The popular cultivars in the middle and lower reaches of Yangtze River were derived from wild populations of *Z. latifolia* in the same area. The aim of this study was to detect the interaction between the heterogeneous environmental variables and potential molecular basis of local adaptation. By doing so, our specific questions are as follows: (1) What is the genetic structure of *Z. latifolia* along the latitudinal gradient? (2) How do the environmental factors associated with latitudinal regions affect population divergence? (3) Whether *Z. latifolia* exhibits local adaptation, and how is the local adaptation related to genetic variation? The joint study of genotypes, phenotypes and environments of *Z. latifolia* wild populations would help predict the fate of the plant in global climate changing context, and put forward appropriate conservation strategies.

## MATERIALS AND METHODS

### Sample collecting

In the autumn of 2015, *Z. latifolia* wild populations were collecting from five river basins (Heilongjiang River Basin, HLJ; Liaohe River Basin, LHR; Huanghe River Basin, HHR; Yangtze River Basin, YZR; and Zhujiang River Basin, ZJR), which belong to different latitudes (N 20°21′ – 50°54′). The individuals were transplanted in the Wu-han Botanical Garden, Chinese Academy of Sciences (30°34′55′′N, 114°16′05′′E), and later we established a common garden of *Z. latifolia*. In the summer of 2020, a total of 60 individuals were sampled from the common garden, with six individuals per population and two populations from each river basin. Additionally, three cultivar samples were collected from Wuhan Academy of Agricultural Sciences for sequencing. Young fresh leaves were sampled, and immediately frozen in liquid nitrogen and preserved until DNA extraction. Total DNA was extracted using modified CTAB protocol (Murray & Thompson, 1980).

### Library construction

To prepare the restriction site-associated DNA (RAD) library and perform the sequencing, the genomic DNA underwent a 30-µl reaction with the restriction enzyme *Eco*RI, followed by P1 adapter ligation utilizing T4 ligase. The resulting fragments were subsequently sheared and size-selected randomly to retain those that measured from 350-550 bp. Then, P2 adapter ligation ensued, and the ligation products underwent purification and PCR amplification, followed by size selection to 350-550 bp through gel purification. The quality and quantity of the libraries were determined through the use of Agilent 2100 Bioanalyzer and quantitative PCR, respectively. The paired-end reads (150 bp) were sequenced at Novogene Co. Ltd. (Beijing, China) using two lanes of Illumina HiSeq 2000.

### Processing of Illumina data

RAD-seq sequencing data quality was firstly evaluated using FastQC (Andrews, 2010), and low-quality bases with base quality scores below 20 were removed using Trimmomatic (Bolger et al., 2014). Read lengths less than 150 bp were discarded. BWA MEM (Li & Durbin, 2009) was employed to map the remaining reads to the *Z. latifolia* reference genome (Guo et al., 2015) with default parameters. To facilitate SNP calling, the mapping output SAM files were converted into BAM, with read mate pairs fixed, sorted and indexed using Samtools (Li et al., 2009). ANGSD (Korneliussen et al., 2014) was used for SNP calling, which is specifically designed for low depth of coverage data when a reference genome is available. Biallelic SNPs were retained after calling SNPs with filters of minimum mapping quality of 30 and a SNP *P*-value of 1e-6. Vcftools (Danecek et al., 2011) were used to filter low-quality SNPs for genetic structure, EAA, and coalescent simulations. The POPULATIONS module in the STACKS pipeline (Catchen et al., 2013) was utilized to generate downstream population genetic analysis datasets, where polymorphic RAD loci present in 95% of individuals were kept (*r* = 0.95).

### Genetic diversity and population structure

Genetic diversity estimators including nucleotide diversity (*P*i), expected heterozy-gosity (*H*_E_), observed heterozygosity (*H*_O_) and inbreeding coefficients (*F*_IS_) were cal-culated using POPULATIONS module in STACKS with unpruned SNPs.

We used Bayesian clustering and principal coordinate analysis (PCoA) to estimate population structure. For Bayesian clustering, STRUCTURE v.2.3.4 (Pritchard et al., 2000) with the admixture model was used. To ensure that loci were unlinked as expected by STRUCTURE, we pruned the dataset (kept 9389 SNPs) using the indep-pairwise function (1000 100 0.2) in PLINK1.9 (https://www.coggenomics.org/plink/1.9/). To determine the optimal number of groups (*K*), we ran 10 iterations for *K* 1-10. Each iteration was performed for 200 000 Markov chain Monte Carlo (MCMC) generations with a burn-in period of 100 000 generations. The optimal *K* was chosen using the delta-*K* method implemented in STRUCTURE HARVESTER (Earl & vonHoldt, 2012). CLUMPP (Jakobsson & Rosenberg, 2007) was used to average the cluster membership coefficient for each individual across 10 independent runs and clustering map plotted by DISTRUCT (Rosenberg, 2004). Principal coordinate analysis (PCoA) as implemented in DART R v.1.1.11 (*gl.pcoa* function) was used to estimate the distribution of genetic variation.

Due to uneven sampling across our sampling range, Bayesian clustering may tend to partition continuous variation into spurious clusters and overestimate the number of potential clusters (Gao et al., 2021). We used the conStruct software (https://github.com/gbradburd/conStruct) that infers genetic structure by considering the continuous pattern of differentiation in geographically separated populations (Bradburd et al., 2018). Two models, a non-spatial model, which is similar to the model used in STRUCTURE, and a spatial model that allows for testing for the presence of isola-tion-by-distance were evaluated using this software. For both models, 10 repetitions for *K* = 1-10 each run at 50 000 iterations was performed and cross-validation done to determine the most likely value of *K* and the best model for our data. SNPhylo software (Lee at al., 2014) was used to generate the ML tree with 1000 bootstraps for the pruned SNPs.

### Demographic history

Based on the time estimated for divergence between northern and southern clades (∼ 0.0633 mya) using plastome data (unpublished), eight demographic models were designed. Here four models were designed as test models and the other four as control models. Considering that *Z. latifolia* was introduced in China through the north after which it dispersed southwards (Guo & Ge, 2005), the northern clade identified by genetic structure analyses was considered the ancestral population. Model 1-2 and model 3-4 assumed that after splitting at a time back in history, the two groups continued with constant expansion (model 1-2) or exponential expansion (model 3-4), with or without gene flow. Model 5-6 and model 7-8 assume that after splitting, the two groups continued with constant expansion followed by exponential expansion (Model 5-6) or constant expansion followed by a bottleneck event from which the populations in both groups have never recovered from (model 7-8), with or without gene flow. The simulations were performed using fastsimcoal2 (Excoffier & Foll, 2011; Excoffier et al., 2013). We imputed the 0.05 missing data in our VCF file since fastsimcoal2 models with missing data can result in biased estimates of the site frequency spectrum (SFS). A custom python script *easySFS* was used to generate a 2D joint folded SFS for north clade (n = 36) and south clade (n = 30) using raw SNPs (82,235 SNPs) that were neither pruned nor filtered for rare alleles (MAF<0.05). We used a mutation rate per site per year value of 6.5 × 10^−9^ (Guo et al., 2015; Haas, 2021), and generation time of one year for all coalescent simulations, which has been suggested for wild rice. We run 200 000 simulations 100 times for each model to calculate composite likelihoods (Papadopoulou & Knowles, 2015) and Akaike information criterion (AIC) was used to select the best run of each model and the best model that fit our data.

### Environmental association analysis

The role of the environment in shaping the observed genetic structure was assessed using gradient forest (GF: GRADIENT FOREST v.0.1) (Jiang et al., 2019) and redundancy analysis (RDA: *vegan* R package). The non-parametric machine-learning regression approach of the GF provides the means to explore the association of allele frequency, spatial and environmental variables. GF was used to identify the most important environmental variables which explain the allelic change along the latitudinal range. The association between the identified environmental variables was assessed us-ing RDA. Using the same procedure, we also assessed the role of the environment on the morphological variation of the sequenced samples in the common garden.

For the identification of outlier loci, different software have advantages and disadvantages, which will yield slightly different results. In this study, two methods were used to identify outlier loci; including *pcadapt* and Bayenv2 methods. The *pcadapt* was run for *K* =1-10 with default parameters. Outliers loci were selected after correction using the *qvalue* method. Bayenv2 (Coop et al., 2010; Günther& Coop, 2013), a modified outlier detection method, was used to supplement *pcadapt*. We generated a neutral covariance matrix to be used as a null model in Bayenv2 by using the pruned SNPs and filtering for loci identified as outliers by *pcadapt*. The analysis was performed for 5 independent runs with 100,000 MCMC with the covariance matrix as the null model for calibration. We used the data-set including 18368 SNPs (without pruning) for these analyses, considering that significant associations are likely to be near the true target loci of selection (Gao et al., 2021). Loci that were identified by *pcadapt* and were found within the top 1% of the Bayes factor and 5% of the Spearman’s rank correlation coefficient in Bayenv2 were considered adaptive loci (Gao et al., 2021).

Adaptive loci that were outliers based on *pcadapt* and significantly associated with environmental variables based on Bayenv2 were extracted from the genome using a custom script in FASTA format. Annotation was performed using the BLAST2GO 6.0.1 (Gotz et al., 2008) by BLASTx against non-redundant *Oryza sativa* protein database filtering out hits with e-value < 1.0E-5.

### Phenotypic trait analysis and genome-wide association analysis

Trait data for all sequenced individuals included plant height, tiller numbers, stem diameter, leaf length, leaf width, leaf area, dry weight, number of internodes, internode length and flowering period collected from our common garden experiment. To examine whether morphological and genetic variables showed similar population differentiation, we performed PCA analysis based on the morphological traits. Furthermore, the difference of traits between the two genetic groups identified by structure analysis was tested using Welch’s two-sample *t*-tests assuming unequal variance (Gao et al., 2021). Trait—environment association analysis was conducted using multiple regression models. Here, predictors were standardized to a mean of zero and a variance of one for direct comparison of coefficients. Forward selection using *ordiR2step* function in *vegan* R package (Oksanen et al., 2016) was used to maximize the degree of freedom of the error considering the few replicates per population. Here, parameters were added into the model provided they decreased the AIC. Furthermore, RDA analysis was performed to investigate the association of morphological variation with geographical and environmental factors.

All SNPs (82,235) were used to conduct genome-wide association analysis for the phenotype variables. Multidimensional scaling (MDS) of the genotype data was done, resulting in five principal components (PCs). Generalized linear model (GML) was used to associate SNPs with each of the morphological variables, while incorporating the five PCs as the covariate to correct for population structure. Significant associations were considered after Bonferroni and false discovery rate adjustment at *P*-values (<0.05). Significant SNPs for each trait were extracted together with 125 bp flanking regions using a custom python script. The putative genes were annotated through BLASTx using *Oryza sativa* in Blast2Go v.6.0.1 (Conesa and Götz, 2008).

### Isolation by distance and isolation by environment analysis

Neutral loci (pruned and non-outlier SNPs), adaptive loci (SNPs identified by *pcadapt* and bayenv2 as outliers and significantly associated with climatic variables) and phenotype from common garden data for the individuals we sequenced, were used to assess for patterns of IBD and IBE. Here, we performed simple, partial and combined RDA to test for influence of IBD, IBE on both allele frequency and morphological variation. Allele frequency matrix was generated from the pruned SNP data and transformed us-ing Hellinger method (Legendre & Gallagher, 2001). Geographic coordinates in decimal degrees and transformed in *vegan* were used to compute the geographical matrix. Significance of each partition was assessed using *anova.cca* function of the *vegan* package at 999 permutations (Oksanen et al., 2016). Environmental variables were also forward selected to retain the most important ones using the *ordiR2step* function in *vegan*. Two independent matrices were used, including the 12 environmental variables (representing IBE) and geographic variables (IBD). RDAs were performed for the neutral and adaptive SNPs, and morphological variation of the samples.

## RESULTS

### RAD-seq, genetic diversity and structure

Sequencing of the RAD library generated 305,747,458 clean reads with an average of 5,095,792 reads per individual. After de-duplexing, a total of 242,590,068 clean reads were retained, with an average of 4,043,167 reads per individual. The RAD sequences covered approximately 2.7% (average 16.74 Mbp, with a range of 14.94 to 19.71 Mbp among individuals) of the *Z. latifolia* genome (Table S1), which is 600 Mbp in size. The average sequencing depth was 3x ranging from 2x to 5x (Table S1), which was considered sufficient to answer our research questions. This is because a quality reference genome and a bioinformatics tool (NGStools) that is designed for working with low coverage RAD data were available.

Low levels of genetic diversity were observed, especially at the species distribution periphery (Table 1). Higher than average diversity was observed along the central region. The inbreeding coefficients (*F*_IS_) ranged from −0.14893 (population CTV) to 0.05673 (population DT), which suggested that none of the populations showed heterozygote deficiency. Extremely low numbers of private alleles were found in the south most populations FC and DC (Table 1).

**Table 1:** Genetic diversity estimates of *Zizania latifolia* populations using all SNPs.

| Group ID | Pop ID | Private alleles | N | $H_O$ | $H_E$ | $P_i$ | $F_{IS}$ |
| --- | --- | --- | --- | --- | --- | --- | --- |
| N1 | HDY | 420 | 6 | 0.132 | 0.087 | 0.096 | -0.070 |
| N2 | YLZ | 411 | 6 | 0.088 | 0.080 | 0.088 | 0.002 |
| N3 | HR | 557 | 6 | 0.133 | 0.099 | 0.108 | -0.049 |
| N1 | DG | 458 | 6 | 0.150 | 0.137 | 0.150 | -0.002 |
| N4 | HXD | 150 | 6 | 0.132 | 0.102 | 0.112 | -0.041 |
| N4 | MK | 228 | 6 | 0.150 | 0.141 | 0.155 | 0.010 |
| S1 | DT | 412 | 6 | 0.141 | 0.152 | 0.166 | 0.056 |
| S1 | BD | 736 | 6 | 0.163 | 0.146 | 0.159 | -0.001 |
| S2 | FC | 10 | 6 | 0.202 | 0.126 | 0.138 | -0.122 |
| S2 | DC | 6 | 6 | 0.196 | 0.125 | 0.137 | -0.112 |
| S3 | CTV | 1279 | 6 | 0.229 | 0.138 | 0.151 | -0.148 |

**Note:** Group ID corresponds to clusters identified by PCA. Pop ID corresponds to the population names except for CTV which stands for cultivated individuals. N represents north group while S represents south group.

Genetic structure analysis showed that the samples clustered into two clades, the north and south clades using STRUCTURE, SNPhylo and PCoA (Figure 1 & 2). However, each clade was further clustered into 4 and 3 sub-groups respectively. The supplementary genetic structure analysis implemented in software conStruct supported the two groups identified by STRUCTURE. Here, the spatial differentiation model showed presence of a weak IBD and divided the total variance into two layers corresponding to the north and south clades (Figure S1).

**Figure 1:**
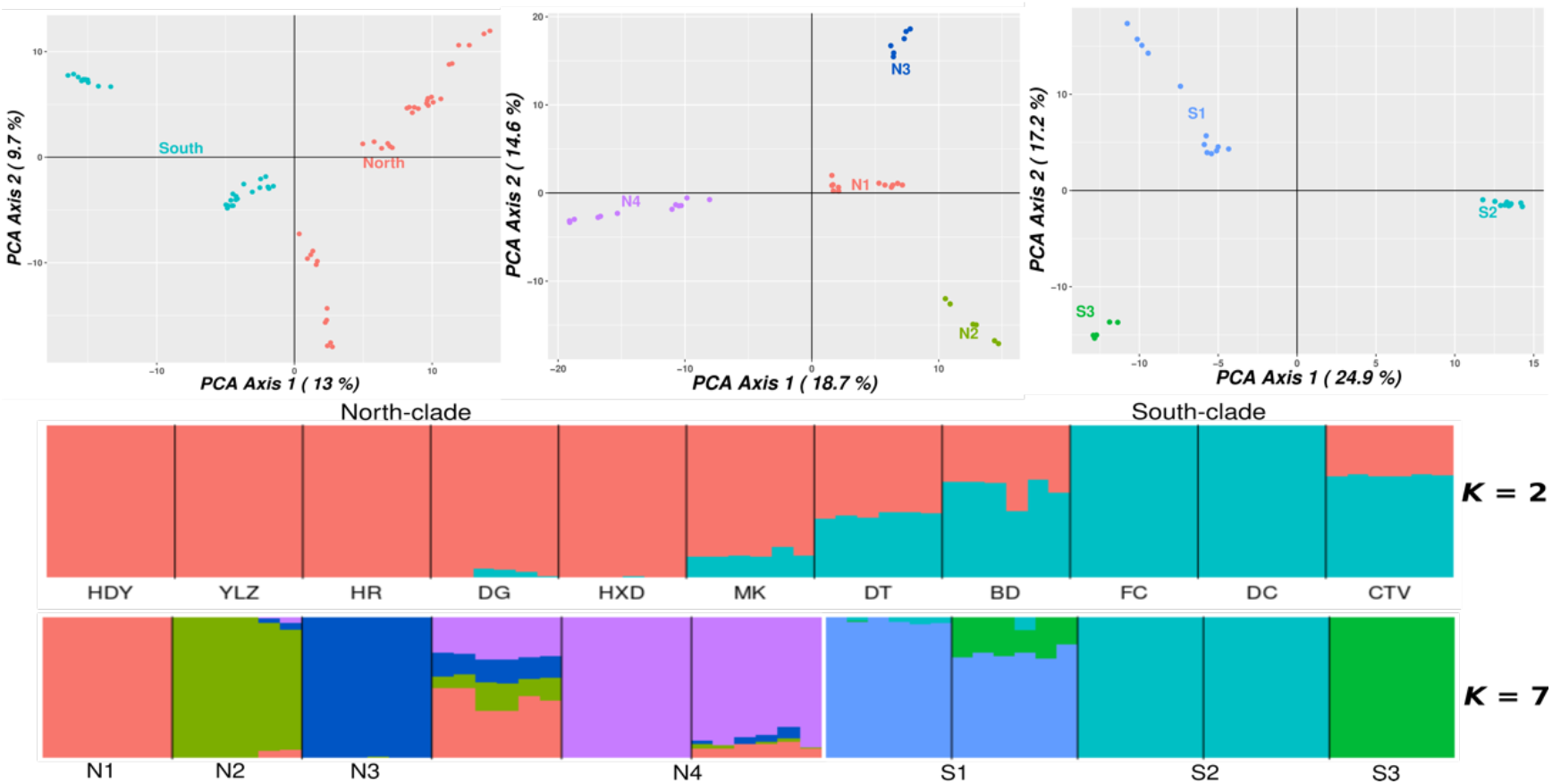
STRUCTURE and principle component analysis (PCoA) showing the clustering of populations into north and south clades and into seven sub-groups based on river basin from which the samples were collected. N1 and N2, Heilongjiang; N3, Liaohe; N4, Huanghe; S1, Yangtze; S2, Zhujiang; S3, cultivated samples from Yangtze region.

**Figure 2:**
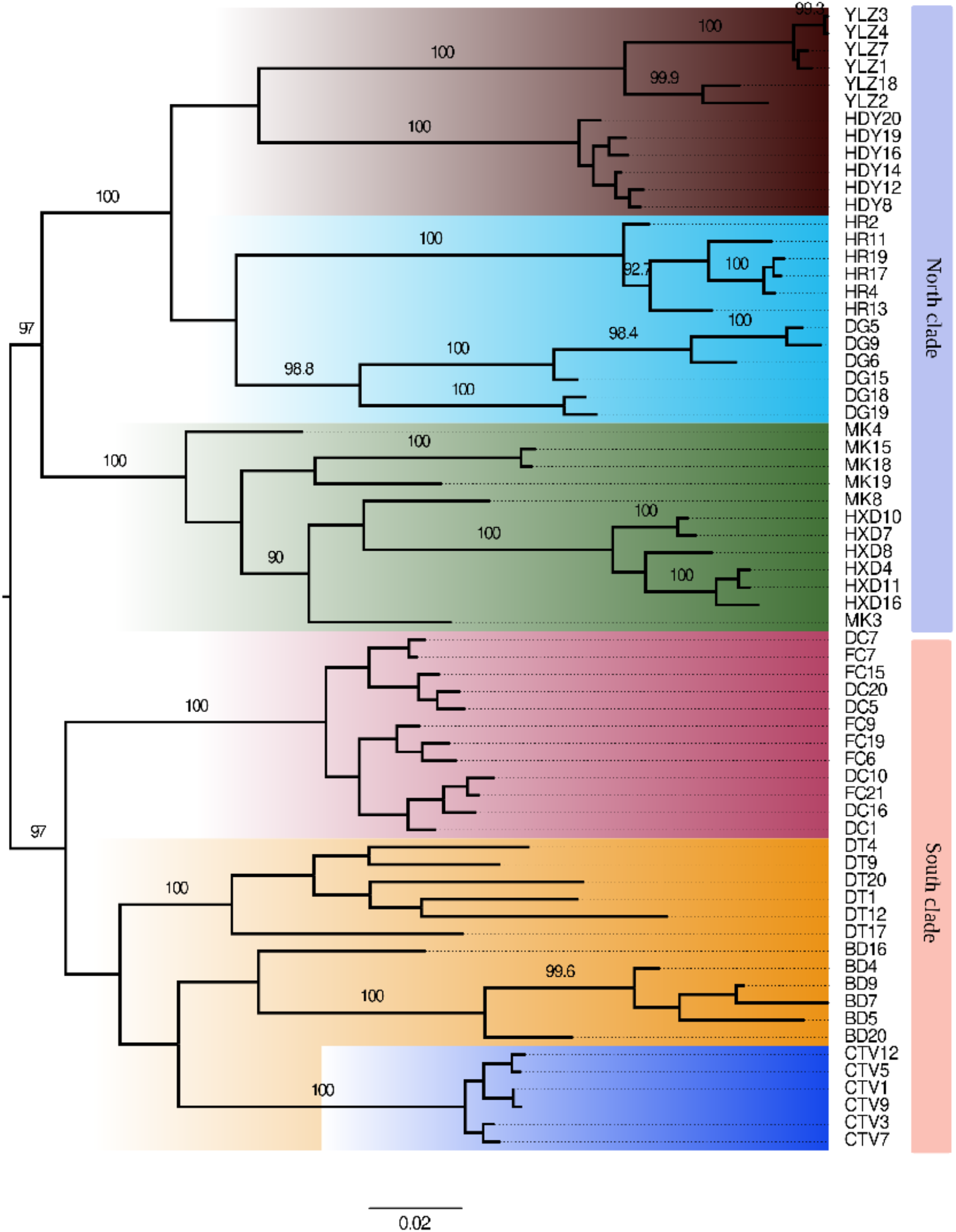
Pruned 9389 SNPs ML tree generated using SNPhylo. The highlight colors correspond to the 5 River Basins from which the samples were collected. Brown, Heilongjiang; light blue, Liaohe; light green, Huanghe; purple, Zhujiang; gold, Yangtze; blue, cultivated samples.

Pairwise *F*_ST_ revealed that populations from the north (HLJ) had the highest differentiation (*F*_ST_ = 0.29), and differentiation decreased southwards reaching the lowest (*F*_ST_ = 0.03) in the southmost population ZJR (Table S2).

### Population history

For the demographic simulations models using fastsimcoal2, we found that model 8, which was simulated for constant expansion with gene flow after splitting of both populations groups followed by a bottleneck event from which the populations have never recovered from, as the best model that fit our data using the AIC. In brief, the north and south clades, after splitting 8k years ago, both expanded constantly with minimal gene flow, and about 2k years later, they both experienced genetic bottlenecks, with no gene flow and they have not recovered to date. Exponential expansion, simulated by model 5-6, was the least supported, which indicated that there was less possibility that the population size for the two groups has been increasing (Figure 3, Figure S2)

**Figure 3:**
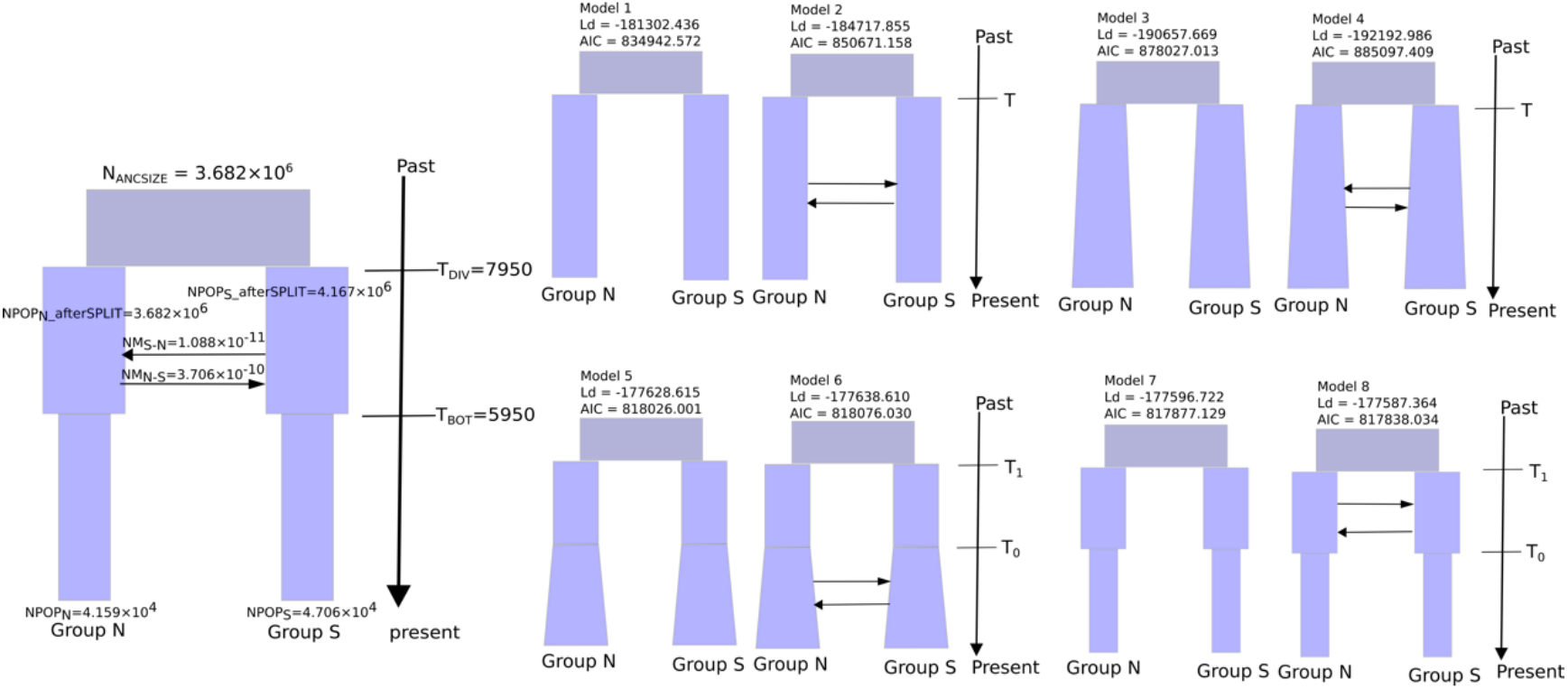
Demographic models simulated using SFS in fastsimcoal2. Model1 and model2, constant expansion after split; model3 and model4, exponential expansion; model5 and model6, constant followed by exponential expansion; model7 and model8, constant followed by bottleneck with or without gene flow.

### IBD and IBE

Environment alone had a higher contribution compared to geographical distance for allele frequency variation for both neutral and adaptive loci, and for morphological variation. Environment, while controlling for geographical distance had also a higher contribution to allele frequency variation and morphological variation compared to geographical distance when controlling for environment. In addition, both environment and geography had the highest contribution in genome variation and trait variation. Most of the genome variation (73% for neutral loci and 90% for adaptive loci) could be explained by both environment and geography, while over 93% of the morphological variation could be explained by both environment and geographical distance (Table 2).

**Table 2:** Contribution of environment and geography in the genome and phenotype variation.

|  | Neutral loci | Adaptive loci |  | Phenotype |
| --- | --- | --- | --- | --- |
| | $r^2$ | $r^2$ | | $r^2$ |
| F~env | 0.4702*** | 0.7545*** | Morp~env | 0.7739*** |
| F~geo | 0.3467*** | 0.6387*** | Morp~geo | 0.611*** |
| F~env\|geo | 0.1713*** | 0.1258*** | Morp~env\|geo | 0.1629*** |
| F~geo\|env | 0.0478*** | 0.0101*** | Morp~geo\|env | 0 |
| F~env+geo | 0.5180*** | 0.7646*** | Morp~env+geo | 0.7739*** |
| Total explained | 0.7371 | 0.9005 | Total explained | 0.9368 |
| Total unexplained | 0.2629 | 0.0995 | Total unexplained | 0.0632 |
| Total | 1 | 1 | Total | 1 |

### Environmental association analysis (EAA)

We retained 10 environmental variables after testing for collinearity using the forward selection method. Together with their first PC, which explained 91.63% of the variation and highly correlated (r>0.6) with bio3, wet-day frequency and UVB, they were then subjected to gradient forest analysis to determine their effect on the allele frequency change along the latitude using pruned SNP data. Temperature, precipitation, UVB2 and wet-day-frequency were the best variables that explained the allele frequency variability along the latitude (Figure S3). RDA analysis showed that the north populations correlated highly with low temperature and high precipitation, while the south populations were correlated with high temperature, high UVB2 and high wet-day-frequency (Figure 4).

**Figure 4:**
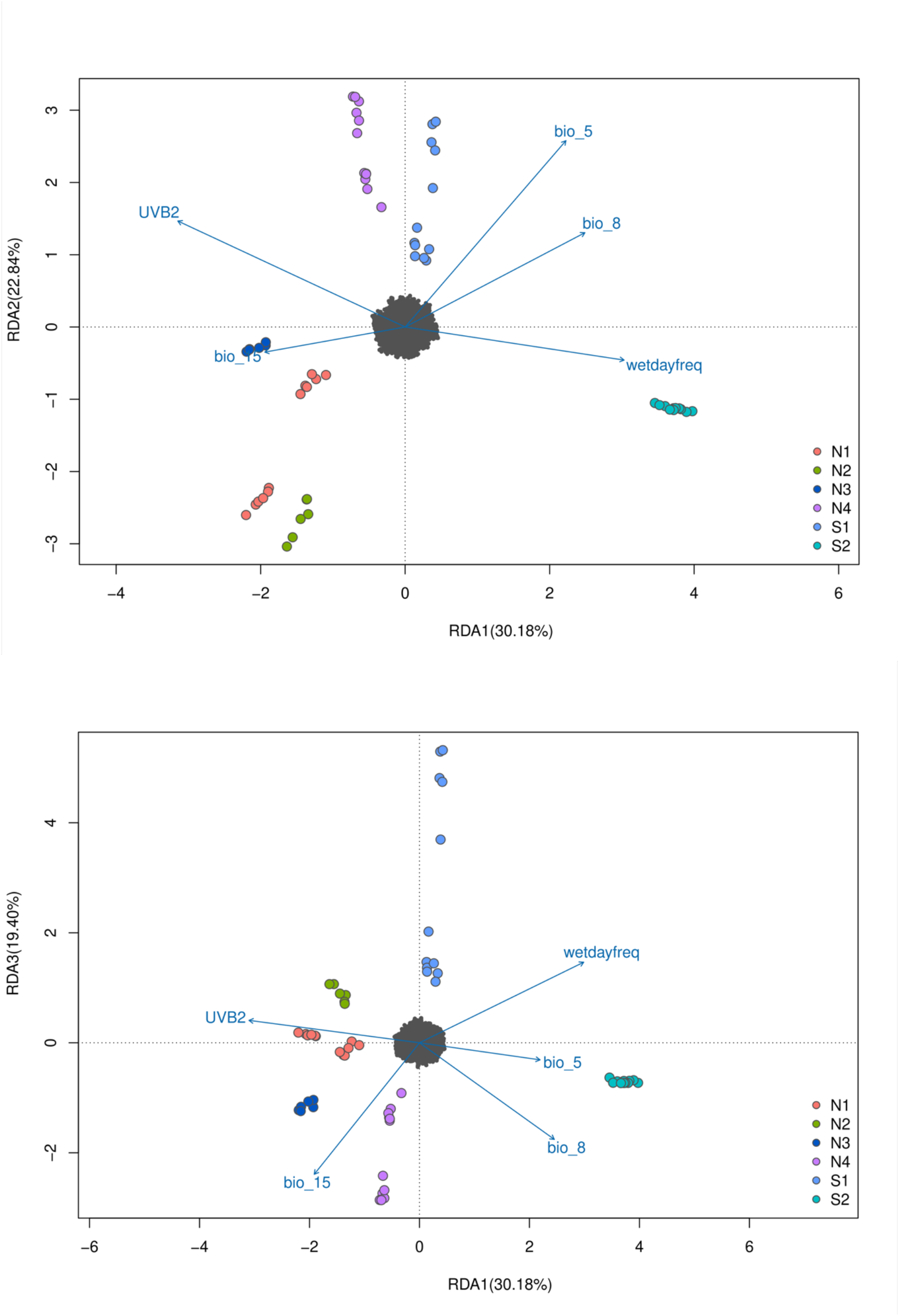
RDA showing the correlation between environmental variables and allele frequency for the groups of populations. N1-N4 represents the north group, while S1-S2 represents the south group.

The similar environmental association analysis performed on morphological variation showed that four environmental factors significantly affected the morpholog-ical variations, including temperature, wet-day-frequency, soil type and elevation (Figure S4). RDA analysis indicated that the southmost populations (DT, BD, FC and DC) are affected by high wet-day-frequency, while the northern populations (HDY, DG, YLZ, HR, HXD and MK) are associated with phaeozems and fluvisols soil type characterized by accumulation of organic matter at the top, low temperature and high elevation (Figure 5).

**Figure 5:**
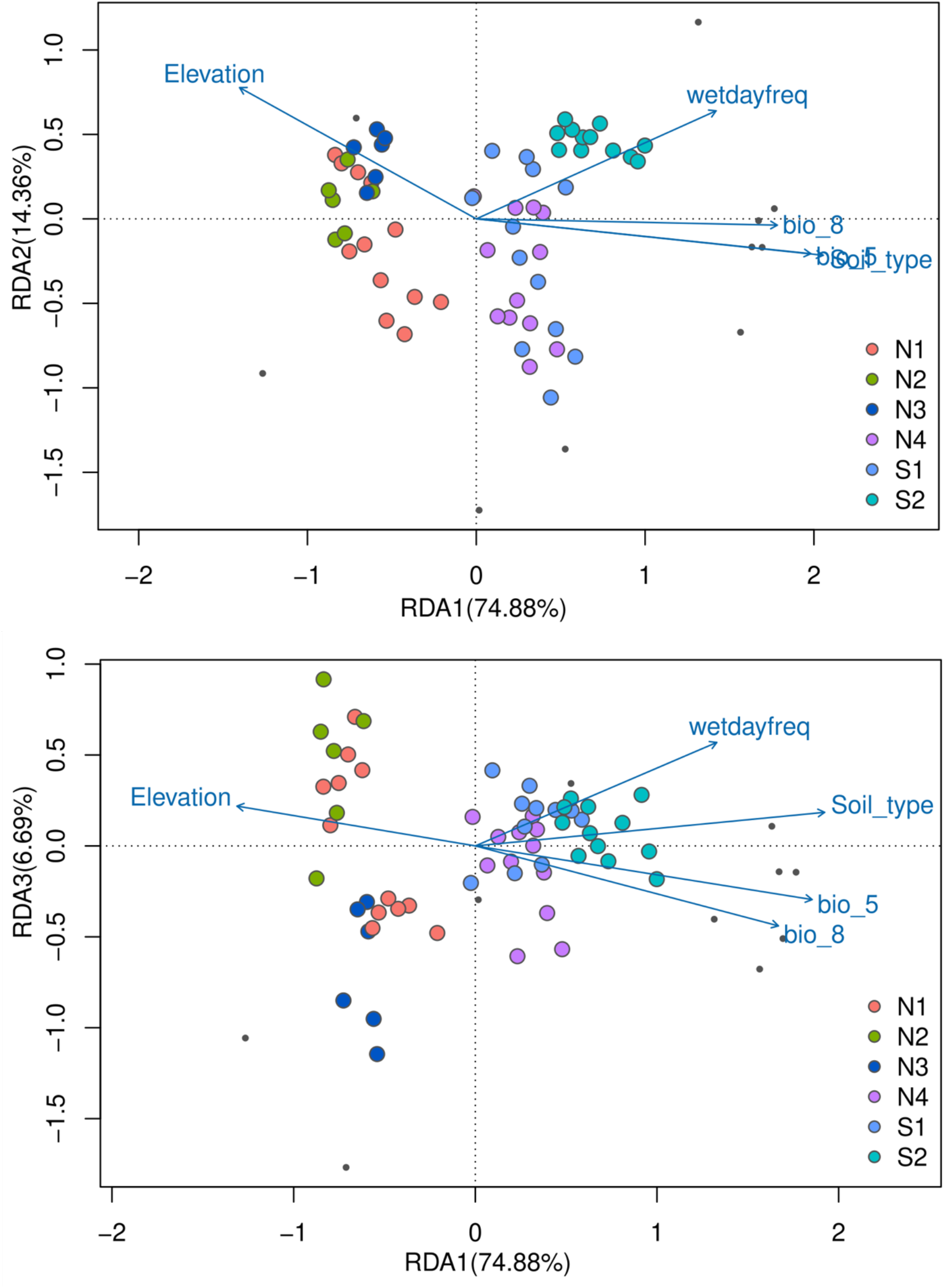
RDA showing the correlation between environmental variables and trait var-iation for the groups of populations. N1-N4 represents the north group while S1-S2 represents the north group.

Using bayenv2 and *pcadapt,* we identified 624 and 1543 outlier loci, respectively. The 281 loci shared by the two methods were considered to be under selection, which highly correlated with PC1, UVB, bio5, elevation and soil type. Annotation of adaptive loci showed that 52 loci were associated with known rice and *Arabidopsis* sequences, majority of which (31 out of the 52) were putative retrotransposons ty1-copia and ty3-gypsy (Table S3). For the remaining sequences, 16 adaptive loci were selected based on the sequence similarity (> 80%) of the genes from rice (*Oryza sativa*) and/or *Arabidopsis* genes. Their putative functions include cell and tissue development, response to UVB, DNA repair, detoxification and defense response (Table 3).

**Table 3:** Putative functions of 16 adaptive loci that had a GO term and >80% similarity BLAST score.

| **Loci** | **Gene name** | **Putative function** | **Length** | **Similarity** |
| --- | --- | --- | --- | --- |
| 104359 | FIM4 | May regulate actin cytoarchitecture, cell cycle, cell division, cell elongation and cytoplasmic tractus | 250 | 97.22 |
| 7075 | Os03g0212700 | Metal ion binding, proteolysis | 250 | 100 |
| 175862 | BBD2 | Involved in basal defense response. Participates in abscisic acid- derived callose deposition following infection by a necrotrophic pathogen | 250 | 94.62 |
| 340982 | NIT4 | May be involved in cyanide detoxification | 250 | 82.05 |
| 119001 | KIN14I | Minus-end microtubule-dependent motor protein involved in the regulation of cell division. | 250 | 100 |
| 281583 | MTR4 | Required for proper rRNA biogenesis and development. | 250 | 100 |
| 177063 | Os03g0294900 | Cellular response to DNA damage stimulus | 250 | 93.02 |
| 192474 | P0560C03.8 | Zinc ion binding, DNA integration | 250 | 86.19 |
| 109853 | WAKL3 | Calcium ion binding, cell surface receptor signaling pathway | 250 | 99.5 |
| 53815 | LOC_Os11g44770 | Metal ion binding, DNA integration | 250 | 90.9 |
| 1448351 | PRL1-IFG | Mediates the ubiquitination and subsequent proteasomal degradation of target proteins | 250 | 80.79 |
| 299511 | LOC_Os11g23750 | DNA integration | 250 | 95.52 |
| 1811459 | Os02g0575700 | Unknown | 250 | 84.49 |
| 729142 | BCHC2 | Nucleotide binding | 250 | 83.98 |
| 235175 | LGD1 | Metal ion binding, cellular amino acid catabolic process | 250 | 83.33 |
| 139162 | TSJT1 | Node development and response to UVB2 | 250 | 84.04 |

### Phenotypic trait analysis and genome-wide association analysis

Among the tested morphological variation, significant difference (*P*<0.05) between the two groups were identified in all traits except for the number of nodes and internode length (Figure S5). The first two principal components accounted for 76.11% of the variation and were consistent with the genetic structure (Figure 6). Plant height, stem diameter, leaf width, area and length, dry weight and flowering were correlated with PC1, while number of nodes and internode length were correlated with PC2.

**Figure 6:**
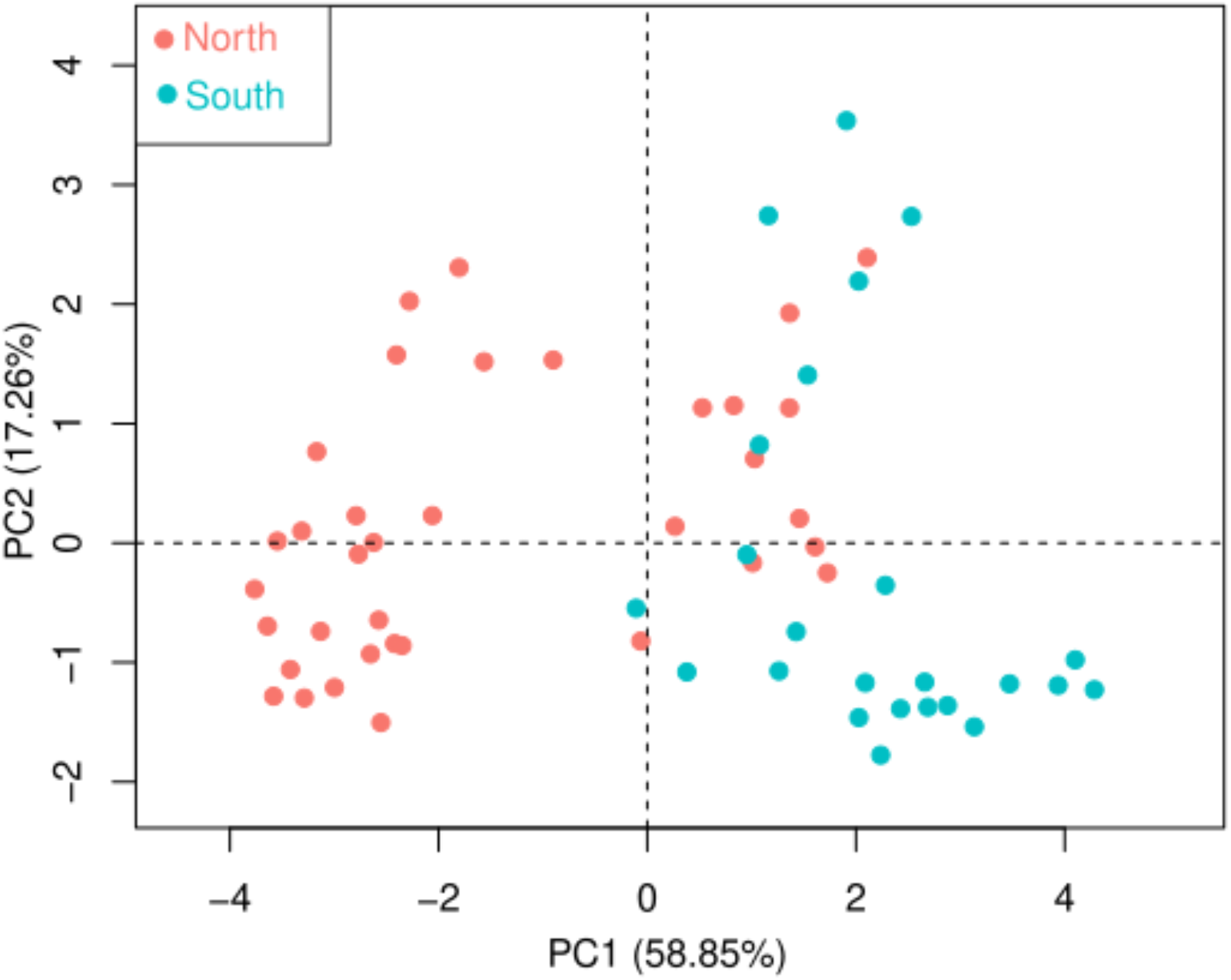
PCA showing morphological differentiation between north clade and south clade of *Zizania latifolia*.

The two population groups inhabit different precipitation and soil pH zones. The northern group was found to be associated with higher precipitation (bio15) (*t*-value 5.43, *P* < 0.05) as well as higher soil pH (*t*-value 2.72, *P* < 0.05). The southern group was found to be associated with higher wet-day frequency (*t*-value −8.31, *P* < 0.05) and clay enriched and human influenced soil-type (*t*-value −3.56, *P* < 0.05) (Table S4).

Different environmental variables showed variation in their effect on different morphological traits (Table 4). Wet-day frequency, precipitation (bio15) and soil-type (anthrosols/human influenced and acrisols/clay enriched) emerged the highest predictors of morphological variation between groups. For example, leaf area, internode length and number of nodes were positively affected by the three environmental variables, while tiller numbers were negatively affected by the same environmental variables. RDA analysis showed that the north populations significantly correlated with low temperature, high elevation and low wet-day frequency, while the south populations were correlated with high wet-day-frequency, high temperature and human influenced soil type (Figure 5).

**Table 4:** Environmental variables that predict variation in measured traits using forward model selection *lm* in R.

| Trait | intercept | Wet dayfreq | elev | bio3 | bio5 | bio8 | bio15 | Soil type | uvb2 | Soil pH | SOC |
| --- | --- | --- | --- | --- | --- | --- | --- | --- | --- | --- | --- |
| Plant height | 0.80 |  | -1.26 |  |  |  |  |  |  |  | -15.64 |
| Tiller number | 127.64 | -20.22 |  |  |  |  | -11.87 | -18.61 | -126.9 |  |  |
| Stem diameter | 0.01 |  |  | 3.62 |  |  |  | 5.10 |  | 8.95 |  |
| Leaf length | 0.42 | -7.17 |  | 31.69 |  | -13.78 |  | 8.54 |  |  | 28.09 |
| Leaf width | -0.59 | 28.39 | 1.68 |  |  | -5.26 | -19.15 | -9.49 |  | 369.55 | -190.85 |
| Leaf area | 0.03 | 15.49 |  | 2.45 |  | -6.89 | 3.54 | 3.53 |  | 69.23 | -49.63 |
| Dry weight | -0.01 |  | 0.05 |  |  |  |  | 0.42 |  | 1.84 |  |
| Nodes number | -21.79 | 14.94 |  |  | -11.08 |  | 5.30 | 2.89 | 22.22 |  |  |
| Internode length | -217.39 | 217.91 |  |  |  | 20.35 | 22.62 | 4.04 |  | -301.69 | 136.95 |
| Flowering | 0.13 | -2.83 |  |  |  | -4.78 | 5.91 | -0.38 |  |  |  |

For GWAS, plant height and internode length traits had the highest number of significantly correlated SNPs, followed by flowering and leaf width. The rest of the measured morphological traits did not correlate with any SNPs (Figure 7). Majority of the loci associated with morphological variation are LTR retrotransposons. We annotated 40 loci that were associated with the four morphological traits. Most of the loci code for putative genes involved in metal ion binding, DNA integration, CHO metabolism, stress response, photosynthesis, water transport, protein phosphorylation and cellular detoxification (Table 5).

**Figure 7:**
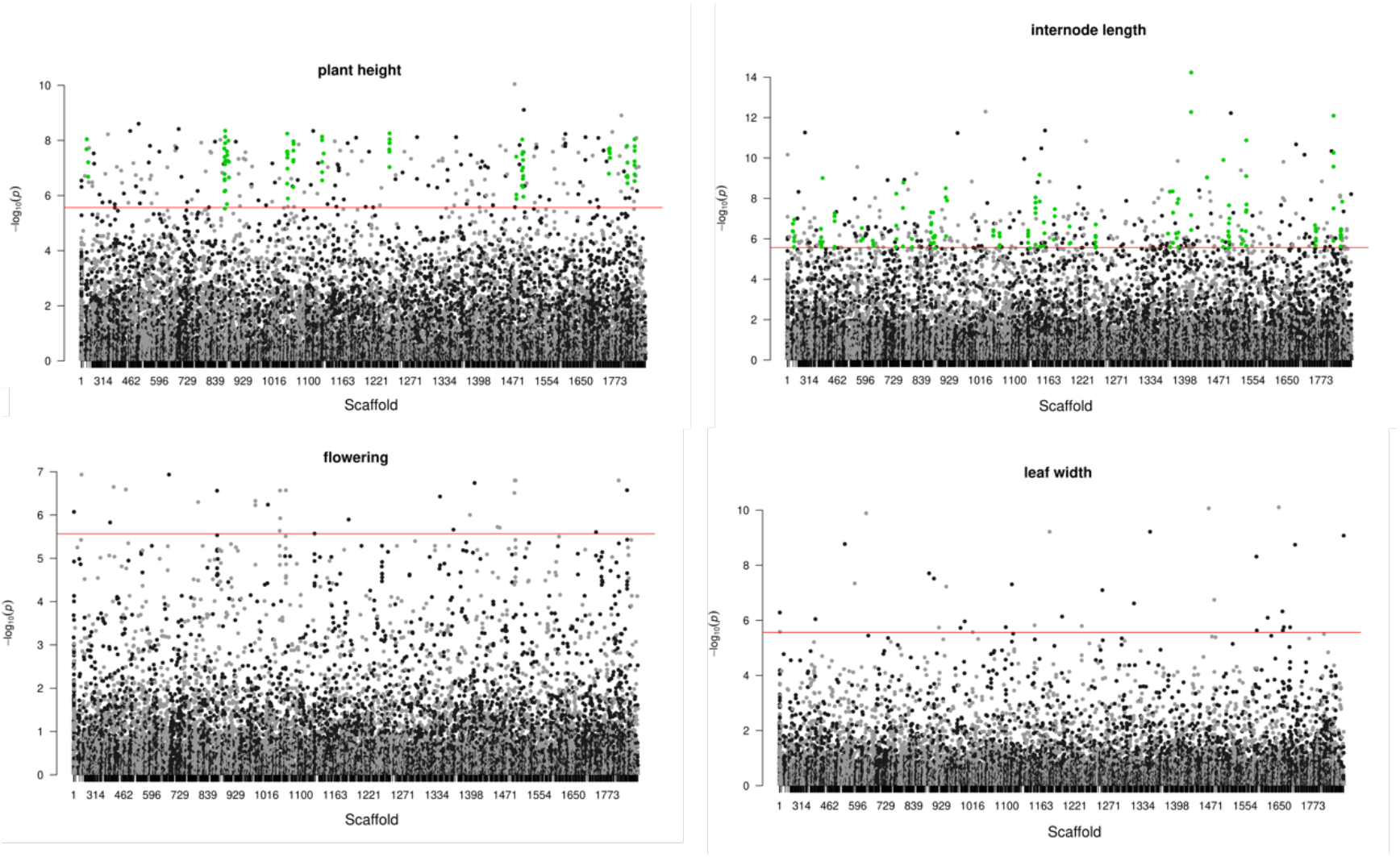
Manhattan plots showing the four traits that were associated with loci in ge-nome-wide association analysis using Tassel5. The green dots represent the regions of the genome that had more than 10 SNPs associated with the trait.

**Table 5:**
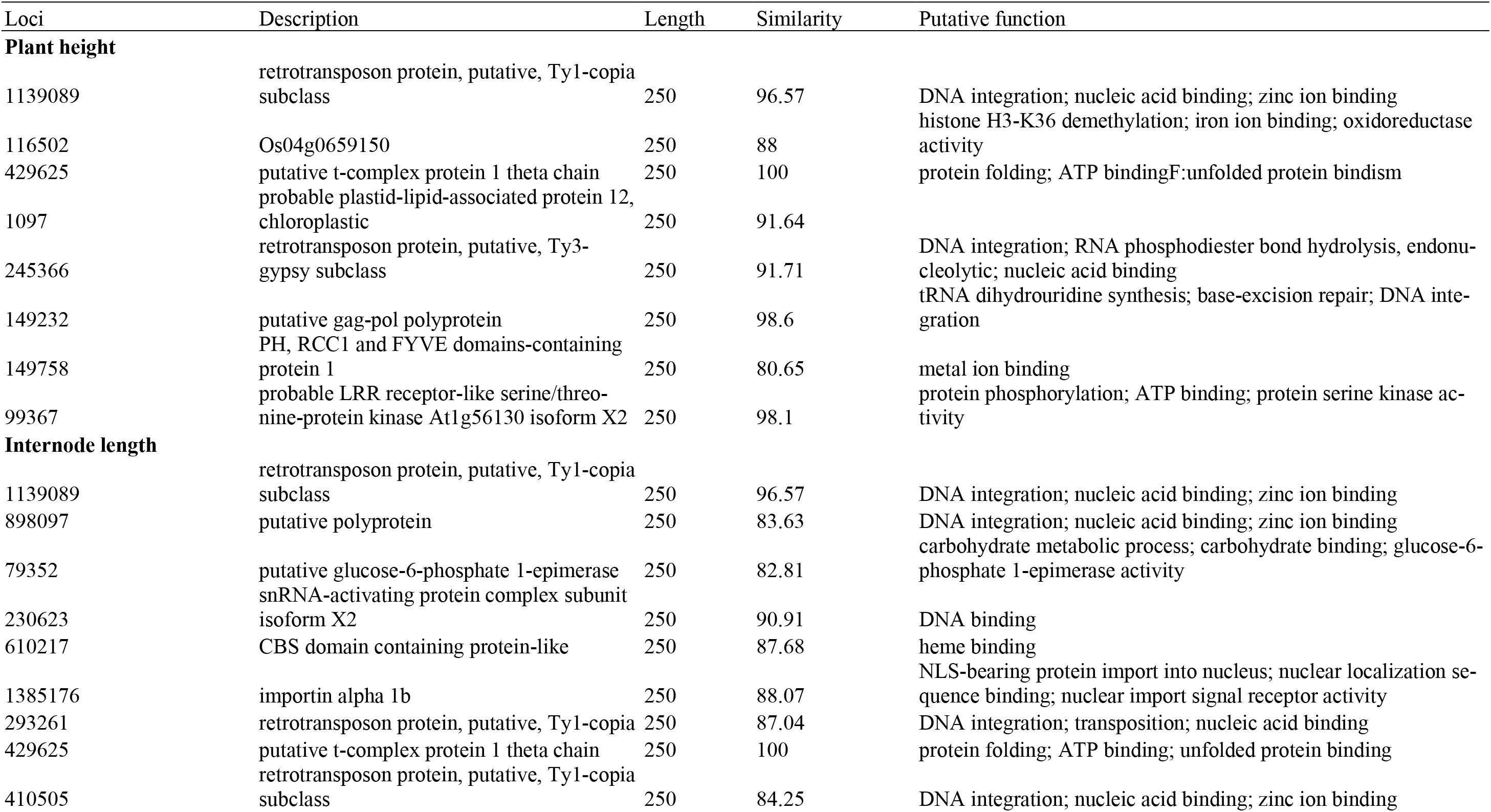
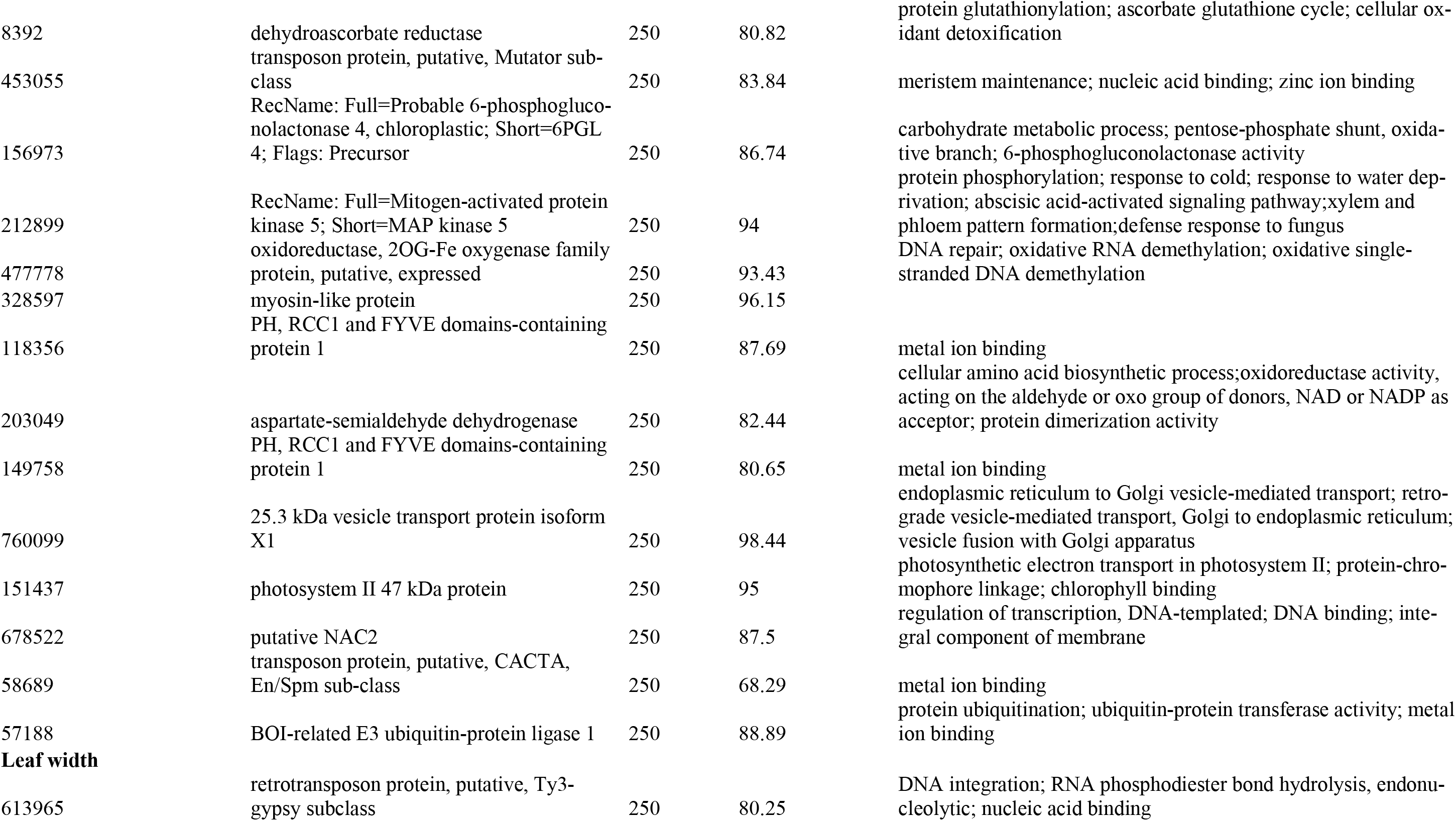
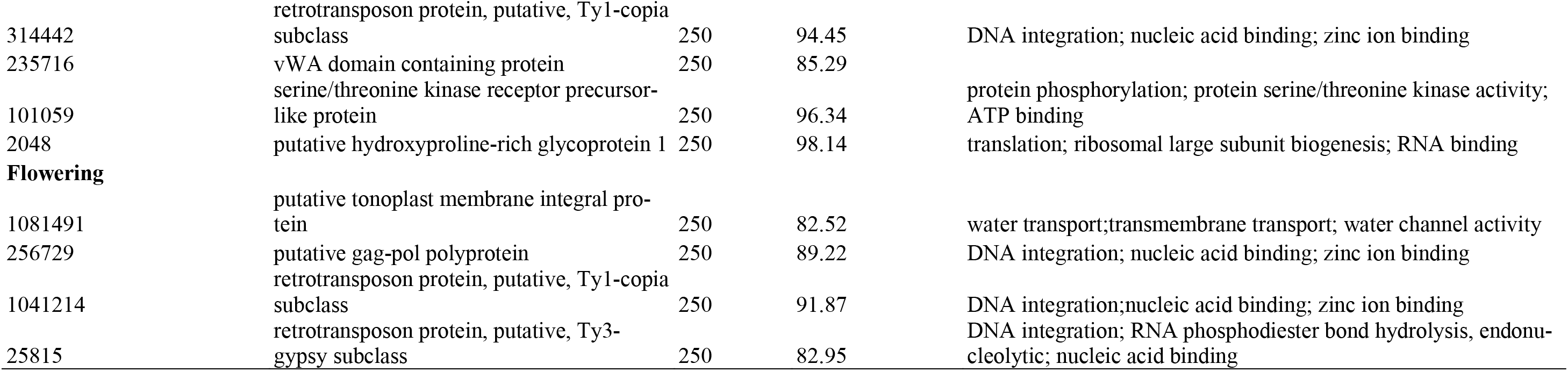
The putative function of 40 loci that were significantly associated with the four traits.

## DISCUSSION

### Genetic diversity

For the 10 *Z. latifolia* wild populations, the highest level of genetic diversity was found in the populations from the Yangtze River Basin in central China (*H*_E_= 0.146-0.152), which was also evidenced by the recent study based on SSRs (Wagutu et al., 2022; Chen et al., 2017). The Yangtze River Basin is the largest alluvial plain in China. In this area, thousands of shallow lakes have connected with the mainstream of the Yangtze River, which forms a stable and extensive potamo-lacustrine habitat. *Zizania latifolia* populations often stretch across the shores of these lakes. Therefore, we inferred that the high level of genetic variation in the Yangtze River Basin might be attributed to the stable habitat and large-scale distribution of the populations. On the contrary, the lowest level of genetic variation was found in the populations from the northernmost Heilongjiang Basin (*H*_E_= 0.080-0.087), which may have resulted from the prolonged hard winters and ephemeral habitats of ponds and ditches.

This trend is characteristic of a combined central-marginal (C-M) and latitudinal pattern of widespread species’ genetic diversity (Guo, 2012). The C-M posits that populations at the center benefit from symmetrical gene flow, suitable climatic conditions compared to the harsh periphery especially in a linear N – S distribution pattern. On the other hand, the latitudinal trend predicts that populations at the lower latitude should show higher genetic diversity that decreases along the latitude due to increased evolutionary stability around the tropics compared to the cold temperate zones. When these two trends are observed for a species, a pattern of higher diversity at the center and low diversity at the periphery, but those at lower latitudinal limits having higher genetic diversity than those at higher latitudinal limits, is expected (Guo, 2012). In this study, the populations from southernmost Zhujiang River Basin exhibited the second lowest genetic diversity, only higher than that of the northernmost Heilongjiang River Basin.

We found extremely few private alleles (only 6-10) in the populations from the Zhujiang River Basin compared to the rest of the populations (Table 1). Such low numbers of private alleles have been attributed to genetic similarity within populations, mostly due to isolation and inbreeding (Kalinowski, 2004, Wagutu et al., 2020). Isolation and excessive human interference in this region have been cited as the cause of low genetic diversity and differentiation (Wagutu et al., 2022), which could be attributed to the observed low number of private alleles.

Compared with wild populations, cultivated populations often harbor less genetic variation due to genetic bottlenecks because of the limited number of progenitors and subsequent artificial selection (Makino et al., 2018), which was also evidenced by the present study. In this study, the cultivated population from the Yangtze River Basin showed lower level of genetic variations (*H*_E_= 0.138; *P*_i_ = 0.151) than the two wild populations from the same area (DT: *H*_E_= 0.152; *P*_i_ = 0.166 and BD: *H*_E_= 0.146; *P*_i_ = 0.159). It was noted that the cultivated population had the highest number of private alleles (1276) compared to that of the wild populations (6-736), which might be explained by the fact that the cultivated population contained three popular cultivars with distinctive horticultural features, like different yields and maturities.

### Genetic structure

The three approaches (Structure, ML tree and PCoA) consistently showed that all populations were firstly clustered into two clades, the north group and south group, which was also manifested by the recent study based on SSRs (Wagutu et al., 2022). Addi-tionally, the SSRs study further showed that all the individuals from the same population clustered together (Wagutu et al., 2022). This was currently supported by Structure and ML tree, except the individuals from the Zhujiang River Basin. The divergence may result from the fact that SSRs are effective to detect structure at a relatively smaller spatial scale compared with SNP mutations (Tsykun et al., 2017). Interestingly, the cultivated individuals were nested against the population from the Yangtze River Basin and did not form a unique clade as it could have been expected. This indicates that the three common cultivars in this study could have originated from the middle and lower reaches of the Yangtze River.

### Population history

The RDA analysis showed the environmental factors had a higher contribution compared to geographical distance for allele frequency variation, suggesting that IBE better explained the genetic patterns of *Z. latifolia* although weak IBD was also evident. This result was also supported by the recent SSR study of the species from the similar distribution area (Wagutu et al., 2022). The boundary between the north and south groups is located between the temperate zone and subtropical zone. The two areas are characterized by obvious climatic differences, such as wet day frequency, mean temperature of the warmest quarter (bio5), precipitation seasonality (bio15), soil type and soil Ph (Table S4).

It is also worth noting that wild rice was traditionally used as a cereal crop at a time when most natural wetlands were intact, population size of the species was high and radiation southwards was ongoing. However, extensive wetland reclamation, degradation and fragmentation has occurred in the last several decades. This has resulted in population size decline and isolation. Coupled with climatic differences that are gradually increasing due to the ongoing climate change, the north – south split and isolation has been exacerbated. Taking into account these pieces of information, the simulated 8 demographic models showed that the best scenario is where N – S populations expanded constantly after splitting 8k years ago and started to experience bottleneck events 6k years ago from which they have never recovered from (Figure 3). This observation coincides with the 8.2-kiloyear cooling event, characterized by a rapid drop in temperature and changes in precipitation patterns. As the temperature declined in the north, the southern areas relatively remained warmer and habitable, facilitating the migration of species through dispersal agents such as humans, animals and birds. The period between 6k and 7k years ago represents the Holocene climatic optimum that was characterized by humid subtropical climate, which facilitated advancement of farming (Wang, 2021), coinciding with the observed bottleneck in our data. It is also evident from these results that the most unlikely event was exponential expansion after N – S split at any time in history to date. It is thus evident *Z. latifolia* population size has been reducing rapidly and climate difference, which will only get worse with climate change and ongoing human activities within the wetland ecosystems, will continue to impact on the species and that population extinction especially at the distribution periphery could occur sooner than later.

In the previous analysis of the species using SSR markers, we found that IBE was the best model that could explain the observed pattern of differentiation along the eastern regions and most of the river basins, while IBD was the second-best model (Wagutu et al. 2022). Using both adaptive loci and neutral loci as well as morphological data from a common garden experiment, we found that IBE alone and when controlling for IBD was the best model explaining the genetic differentiation of the species. However, the combined effect of IBD and IBE explained most of the genetic and morphological variation (Table 2). As previously discussed, climatic differences along eastern China have been implicated in divergence of populations of species. While the overall eastern China landscape is heterogeneous, the most important features that are known to drive *Z. latifolia* dispersal are wind and water connectivity, both of which are effective at a local scale (Tero et al., 2003). The large latitudinal range, which divided eastern China into approximately eight Eco-geographical zones (Wu et al., 2003), creates unique niches for the species (Wagutu et al., 2022). This climatic isolation coupled with the aforementioned factors such as wetland fragmentation and climate change are the basis for the observed pattern of IBE.

### Signatures of local adaptation

We used a common garden experiment combined with genome-wide association analysis and environmental association analysis to provide insights into local adaptation of *Z. latifolia*. This is because the identified pattern of IBE may not necessarily indicate the presence of local adaptation because the SNPs analyzed using RAD sequences could be anywhere within the genome and their effect on physiological processes that lead to adaptation are unknown (Lowry et al., 2017). Common garden allows for the differentiation of phenotypic plasticity that could be due to non-heritable response to local climatic conditions, and heritable positive selection due to certain environmental conditions (De Villemereuil et al., 2016). As such we found that while, *Z. latifolia* morphological traits (plant height, leaf length, leaf width) among all the populations sampled were basically similar in the field, it was not the case in the common garden.

In the common garden (located in Wuhan, a city from the Yangtze River Basin), the northern populations grew smaller compared to the southern populations. The south group identified by the SNP data comprise populations from Yangtze River Basin and Zhujiang River Basin found in the subtropical and tropical zones in the central and south China, respectively, while the north group comprises the Heilongjiang River Basin, Liaohe River Basin and Huanghe River Basin, which are located in the temperate zones (upper central and north China). The two groups occupy unique niches characterized by significant differences in precipitation, temperature and soil pH (Table S4). Considering the location of the common garden, the Zhujiang River Basin populations were moved up the latitude gradient to a less warm climate, while the northern populations were moved to a much warmer climate.

We observed trade-off responses in the common garden for the northern and southern populations commonly referred to as a flight and fight response, respectively. These responses are characterized by trade-off among flowering/total clonality, vegetative growth and multi-year survival (Wright et al., 2022). Most individuals from the northern populations died after a few years in the common garden, while populations from central and south China were still thriving at the time of writing this paper. The flowering observed in the northern populations, unlike in southern populations, could be a trade-off for multi-year survival, where most resources that could have been allocated to vegetative growth and clonality were channeled towards sexual reproduction in the attempt to increase chances of cross-pollination with native populations and thus better chance of offspring acquiring adaptive traits to thrive in the warmer climate (flight response).

On the other hand, the lack of flowering and superior vegetative growth in southern populations compared to northern populations was interpreted as a fight response. Since the southern populations are adapted to the warm climate, moving them to central China reduced the environmental stress. Thus, they were no longer desperate for their survival and hence sexual reproduction resources were channeled towards rapid clonal growth and spread to colonize the habitat and compete effectively with other species. Taking these results together, it is clear that temperature differences between cooler North China and warmer South China have resulted in adaptive differentiation and local adaptation of *Z. latifolia*.

For the genetic – environment (G – E) association, temperature (bio5, bio8), precipitation (bio15), wet-day frequency and UVB2 were found to best explain the allele frequency change along the latitude (Figure S3). The north clade populations correlated strongly with high precipitation and high UVB2, while south clade populations with high temperature and higher wet-day frequency. The north clade is located within the temperate zones, while the south clade is located in the subtropical and tropical zones, which differ significantly in temperature and precipitation (Table S4). Moreover, UVB2 is strong in lower latitudes and high altitude while wet-day frequency is associated with high temperatures. In fact, RDA analysis of morphological variation found that two variables, temperature and wet-day frequency were important in shaping both allele frequency and morphological variation along the latitudinal gradient (Figure S3, S4). This explains the congruence in the morphological and genetic differentiation that we found using the PCA analyses (Figure 1, 7).

Outlier loci analysis and association with climatic variables identified 281 loci that are putatively under selection. Annotation of these loci found 52 loci were associated with known rice and *Arabidopsis* sequences. Out of the 52 loci, 31 were annotated as retrotransposons ty1-copia and ty3-gypsy (Table S3). Long terminal repeats (LTR) retrotransposons are abundant in plant genomes and are found in a high number of copies (Vicient & Schulman, 2002). For the *Z. latifolia* genome, LTR retrotransposons makeup 29.8% of the genome (Guo et al., 2015). It has widely been reported that although these retrotransposons could be quiescent, stress such as pathogen attack, wounding and climatic cues can activate them and since they are easily copied and integrated elsewhere within the genome, they can greatly increase their copy number and genome size. They have been shown to alter gene expression thus leading to diversification between individuals, populations and species (Haas et al., 2021; Galindo-González et al., 2016; Kalendar et al., 2000). It is thus possible that *Z. latifolia* applies the same mechanisms in order to adapt to the colder north and hot south zones and that the overall genome divergence could partly be due to LTR retrotransposons response to climatic conditions. In fact, the phylogenetic tree using the 281 adaptive loci showed a clear north – south divergence based on the climatic difference, compared to all loci and neutral loci (Figure S6).

Some of the other genes that correlated with climatic variables include TSJT1 gene, which has been found to express in presence of UVB stress and is reported to be involved in internode length elongation. This suggests that the gene inhibits plant growth and development in presence of UVB2, a mechanism that could help the plant decrease the surface area on which it is exposed to UVB. The other gene, Os03g0294900, has been found to be up-regulated in severe water-deficit conditions in rice, suggesting its role in drought tolerance through regulation of embryonic development (Moumeni et al., 2015).

Genome-wide association analysis showed an interesting trend similar to the G – E analysis. Most of the enriched SNPs were annotated as LTR retrotransposons, which was the case with annotation of the adaptive SNPs. The morphological traits that correlated with most of the SNPs include plant height, leaf with, flowering and inter-node length. Growth and development genes were found to be associated with climatic variables in the G – E analysis and especially TSJT1 and Os03g0294900. These genes work together with others unique to different morphological traits such as 2OG-Fe and MAP kinase 5 involved in DNA repair, CHO metabolism, stress response, photosynthesis, water transport, protein phosphorylation and cellular detoxification.

Taken together, the local adaptation in *Z. latifolia* is driven by difference in climatic conditions and mainly temperature between the subtropical and tropical cen-tral-south zone and the temperate north. Heritable genetic differences are majorly through activation of LTR retrotransposons, integration along the genome, their putative role in gene expression regulation and subsequent phenotypic diversity that allows adaptation.

## CONCLUSION

Local adaptation is a complex phenomenon to unravel mostly because it can be shrouded by phenotypic plasticity. Moreover, most genomics studies that study local adaptation tend to perform genome scans to link genotypes with the environment, disregarding phenotypic traits that could potentially be under selection. Phenotypic traits are usually polygenic and detecting the actual loci linked to phenotypic variability from a genome wide scan requires high density markers. Moreover, the genetic basis of adaptation may not necessarily entirely be due to gene evolution, but stable and heritable epigenetic changes as well as action of non-coding sequences such as LTR retrotransposons.

In this study, we have approached the study of local adaptation by jointly studying genotypes, phenotypes and environments of a widespread species. Our results show that various climatic factors such as temperature, precipitation and UVB have significantly led to adaptive differentiation of *Z. latifolia* through activation of retrotransposons, gene evolution and subsequent phenotypic variation. Interesting pattern of flowering adaptation at higher temperature, growth traits (plant height, leaf area) at cooler temperature and higher precipitation – traits that are significantly reduced after individuals are transplanted to different environments – provide an outlook of how the species would behave in the predicted warmer future. We also affirm our previous observation of the existence of local adaptation and the genetic basis of phenotypic plasticity in *Z. latifolia* (Wagutu et al., 2022, Jiang et al., 2023). The central China populations showed relatively better genetic health compared to north and south groups. A similar population fitness pattern was identified by our previous SSR and trait analysis, which necessitates both *in-situ* and *ex-situ* conservation of Yangtze River Basin populations as a priority, but also the north and south germplasms, to increase genetic diversity.

The reported adaptive morphological traits and functional sequences as well as the overall populations adaptive profiles, could aid in designing of conservation strategies, prediction of the species performance in changed future climate and in marker assisted breeding. While the reported functional sequences account for a small part of the identified adaptive loci that were available in annotations databases, further testing of these ecological loci could be done using knockout mutants to increase their usefulness in breeding. Considering the over-representation of retrotransposons in adaptive loci, analysis of gene expression could confirm the correlation of these adaptive genes and the role of transposable elements as gene expression regulators in driving local adaptation.

## DATA AVAILABILITY STATEMENT

The datasets presented in this study can be found in online repositories. The names of the repository/repositories and accession number(s) can be found in the article Supplementary Material.

## AUTHOR CONTRIBUTIONS

XF, YC, and WL designed the study. XF and YC collected populations in the field. GKW, XF, YC, MCT, HKN maintained samples in the common garden. GKW, XF, YC performed the experiments. GKW analyzed the data and together with YC wrote the manuscript. All authors read and approved the final manuscript.

## ACKNOWLEDGEMENT/FUNDING

This work was supported by the Strategic Priority Research Program of Chinese Academy of Sciences (grant number XDB31000000), by the Youth Innovation Promotion Association CAS (2021340), and the CAS-TWAS President’s Fellowship for Interna-tional Doctoral Students.

## Notes

### Competing Interest Statement

The authors have declared no competing interest.

