## Supplemental Materials for "RAD-seq unravels local adaptation along a latitudinal gradient for the wild rice *Zizania latifolia*"

Supplementary Material

Table S1: Sequencing details, genome coverage and mean depth for the samples used in this study

| Sample | Effective Rate(%) | Q30(%) | Clean reads | Deduplicated clean reads | coverage | mean depth |
| --- | --- | --- | --- | --- | --- | --- |
| HDY16 | 97.39 | 92.11 | 6499590 | 5036338 | 16.347 | 2.9721 |
| HDY19 | 97.36 | 92.05 | 4588530 | 3627887 | 17.121 | 1.82431 |
| HDY20 | 97.96 | 91.91 | 4479000 | 3486413 | 17.28 | 2.10004 |
| HDY12 | 97.27 | 92.3 | 5111249 | 4041481 | 16.566 | 2.64442 |
| HDY8 | 97.86 | 91.47 | 3087111 | 2405372 | 14.939 | 2.51158 |
| HDY14 | 98.15 | 91.66 | 4943457 | 3875040 | 15.87 | 4.50643 |
| YLZ1 | 97.91 | 91.53 | 5778160 | 4444042 | 17.284 | 5.15831 |
| YLZ3 | 97.21 | 91.84 | 6537978 | 4928306 | 15.536 | 5.9082 |
| YLZ4 | 97.26 | 91.28 | 4488050 | 3516641 | 16.256 | 2.46419 |
| YLZ7 | 97.78 | 91.41 | 5642709 | 4261174 | 16.923 | 3.03456 |
| YLZ2 | 97.74 | 91.42 | 4659279 | 3590785 | 16.329 | 2.70573 |
| YLZ18 | 97.49 | 91.28 | 5249114 | 4125469 | 17.22 | 2.98697 |
| HR4 | 92.67 | 91.2 | 3903632 | 3129059 | 16.711 | 2.13052 |
| HR17 | 97.32 | 91.54 | 6375711 | 4921075 | 16.273 | 3.43 |
| HR13 | 97.5 | 90.97 | 4735443 | 3776911 | 16.55 | 2.38302 |
| HR2 | 97.62 | 91.41 | 4635505 | 3722933 | 15.197 | 2.4983 |
| HR11 | 96.99 | 92.11 | 5641067 | 4338471 | 16.81 | 3.334 |
| HR19 | 94.75 | 92.17 | 5674279 | 4407398 | 16.6 | 3.50935 |
| DG9 | 96.93 | 91.87 | 5081852 | 3946369 | 15.906 | 3.15238 |
| DG5 | 96.96 | 92.03 | 4438054 | 3494792 | 18.529 | 2.71147 |
| DG18 | 96.84 | 92.12 | 5603519 | 4394693 | 17.117 | 3.57375 |
| DG19 | 96.97 | 92.11 | 3740912 | 2993406 | 17.294 | 2.1951 |
| DG6 | 97.22 | 91.98 | 4554852 | 3563274 | 15.929 | 2.71322 |
| DG15 | 97.43 | 92.03 | 5635397 | 4358910 | 17.742 | 3.38488 |
| HXD7 | 98.26 | 90.91 | 4769697 | 3899782 | 17.051 | 2.86884 |
| HXD11 | 98.44 | 91.13 | 6304321 | 5029631 | 16.828 | 3.81122 |
| HXD16 | 98.07 | 90.79 | 5725778 | 4596276 | 16.504 | 3.35735 |
| HXD8 | 98.06 | 90.68 | 4542405 | 3735007 | 16.207 | 2.87967 |
| HXD10 | 97.04 | 90.17 | 5127301 | 4282311 | 17.2 | 2.7567 |
| HXD4 | 97.3 | 90.06 | 6244736 | 5097196 | 16.362 | 3.39582 |
| MK8 | 97.31 | 90.14 | 5712920 | 4778528 | 17.2 | 3.17761 |
| MK18 | 97.69 | 90.39 | 4875433 | 4011738 | 16.526 | 2.65402 |
| MK19 | 98.1 | 91.53 | 4043055 | 3200924 | 16.4 | 2.52039 |
| MK3 | 98.09 | 91.46 | 5428654 | 4242611 | 15.546 | 3.47075 |
| MK4 | 97.66 | 91.54 | 3661571 | 2895699 | 16.328 | 2.30401 |
| MK15 | 98.01 | 91.45 | 5593620 | 4310736 | 16.361 | 3.45989 |
| DT1 | 97.34 | 91.98 | 6466050 | 4975611 | 16.525 | 4.59818 |
| DT17 | 97.26 | 92 | 4481440 | 3566377 | 15.094 | 3.14401 |
| DT20 | 98.24 | 91.84 | 5420747 | 4220906 | 15.51 | 3.48306 |
| DT12 | 97.65 | 92.2 | 5701366 | 4497828 | 15.593 | 3.85396 |
| DT4 | 98.21 | 89.03 | 3667317 | 3128442 | 16.808 | 2.40315 |
| DT9 | 98.74 | 89.44 | 5639026 | 4714288 | 15.856 | 3.66337 |
| BD7 | 98.3 | 89.03 | 7503309 | 6245506 | 16.814 | 5.33397 |
| BD9 | 97.76 | 89.69 | 6824255 | 5609542 | 16.242 | 4.76498 |
| BD4 | 97.8 | 91.77 | 4919413 | 3855426 | 17.405 | 2.88771 |
| BD16 | 98.43 | 91.86 | 6124784 | 4672031 | 18.829 | 3.37921 |
| BD5 | 98.21 | 91.81 | 5851600 | 4453392 | 18.698 | 3.456 |
| BD20 | 98.13 | 91.66 | 5116544 | 3995491 | 16.21 | 2.95835 |
| FC19 | 96.29 | 91.54 | 2579783 | 2072993 | 16.259 | 1.93251 |
| FC15 | 98.17 | 91.68 | 5929426 | 4521222 | 16.214 | 4.2744 |
| FC6 | 98.2 | 91.13 | 3838332 | 3032277 | 15.959 | 2.91407 |
| FC21 | 98.64 | 91.69 | 4986808 | 3936230 | 15.586 | 3.88253 |
| FC9 | 97.8 | 91.14 | 5340106 | 4314592 | 16.217 | 3.2583 |
| FC7 | 96.24 | 91.53 | 5102247 | 4167288 | 17.851 | 3.06726 |
| DC10 | 97.36 | 91.03 | 4537533 | 3685151 | 16.181 | 2.71908 |
| DC1 | 97.74 | 90.94 | 4409491 | 3634991 | 17.337 | 2.594 |
| DC7 | 97.23 | 90.82 | 4928516 | 4017044 | 18.093 | 3.05372 |
| DC5 | 97.01 | 90.83 | 3818095 | 3146574 | 18.616 | 2.35612 |
| DC20 | 97.47 | 90.87 | 4909981 | 4001378 | 19.705 | 3.15673 |
| DC16 | 97.77 | 90.88 | 4537458 | 3658810 | 15.902 | 2.75965 |
| Average |  |  | 5095792.8 | 4043167.8 | 16.67 | 3.173 |

Table S2: Pairwise FST for all populations of Z. latifolia

|  | HDY | YLZ | HR | DG | HXD | MK | DT | BD | FC | DC | CTV |
| --- | --- | --- | --- | --- | --- | --- | --- | --- | --- | --- | --- |
| HDY |  | 0.335807 | 0.319488 | 0.225238 | 0.329747 | 0.232 | 0.221933 | 0.269317 | 0.328854 | 0.322985 | 0.333969 |
| YLZ |  |  | 0.327662 | 0.244267 | 0.349512 | 0.249757 | 0.246138 | 0.284762 | 0.344975 | 0.339367 | 0.351987 |
| HR |  |  |  | 0.228466 | 0.324609 | 0.229697 | 0.21814 | 0.25778 | 0.314739 | 0.308374 | 0.316465 |
| DG |  |  |  |  | 0.229425 | 0.158249 | 0.15541 | 0.195398 | 0.240422 | 0.235093 | 0.241927 |
| HXD |  |  |  |  |  | 0.137811 | 0.203182 | 0.242477 | 0.292331 | 0.287081 | 0.303061 |
| MK |  |  |  |  |  |  | 0.134785 | 0.167032 | 0.203227 | 0.198006 | 0.210358 |
| DT |  |  |  |  |  |  |  | 0.135695 | 0.19085 | 0.186064 | 0.183471 |
| BD |  |  |  |  |  |  |  |  | 0.221633 | 0.217307 | 0.204071 |
| FC |  |  |  |  |  |  |  |  |  | 0.0241437 | 0.273107 |
| DC |  |  |  |  |  |  |  |  |  |  | 0.267234 |

Table S3: Adaptive loci associated with known rice and *Arabidopsis* sequences

| Sequence Name | Description | Length | e-Value | Similarity (%) | GO IDs |
| --- | --- | --- | --- | --- | --- |
| KE372114.1_104359 | putative fimbrin | 250 | 8.52008E-05 | 97.22 | P:GO:0051017; P:GO:0051639; F:GO:0000166; F:GO:0003779; F:GO:0005524; F:GO:0051015; C:GO:0005737; C:GO:0005884; C:GO:0032432 |
| KE371097.1_7075 | probable mitochondrial-processing peptidase subunit beta, mitochondrial | 250 | 2.12842E-05 | 100 | P:GO:0006508; F:GO:0004222; F:GO:0046872; C:GO:0005739 |
| KE372901.1_175862 | bifunctional nuclease 1 | 250 | 9.99668E-09 | 94.62 | P:GO:0006464; F:GO:0004518;  C:GO:0005737 |
| KE373027.1_170485 | retrotransposon protein, putative, Ty1-copia subclass | 250 | 8.81209E-09 | 87.6 | P:GO:0006259; F:GO:0003676 |
| KE371119.1_914597 | retrotransposon protein, putative, unclassified | 250 | 9.43594E-11 | 84.68 | P:GO:0006259; F:GO:0003676 |
| KE372063.1_175511 | uncharacterized protein LOC4330109 isoform X1 | 250 | 3.67386E-11 | 94.35 | C:GO:0016020 |
| KE372722.1_282314 | DExH-box ATP-dependent RNA helicase DExH9 | 250 | 4.85636E-12 | 82.22 | P:GO:0006139; P:GO:0009056; F:GO:0000166; F:GO:0003723;  F:GO:0016787 |
| KE372936.1_47809 | retrotransposon protein, putative, Ty3-gypsy subclass | 250 | 3.66434E-12 | 82.18 | P:GO:0006259; F:GO:0003676;  F:GO:0004518 |
| KE370772.1_1284822 | retrotransposon protein, putative, Ty1-copia subclass | 250 | 1.32286E-12 | 82.75 | P:GO:0006259; F:GO:0003676 |
| KE372278.1_340982 | bifunctional nitrilase/nitrile hydratase NIT4-like | 250 | 3.06212E-13 | 82.05 | P:GO:0008152; P:GO:0042221;  F:GO:0016787 |
| KE372048.1_201093 | retrotransposon protein, putative, Ty1-copia subclass | 250 | 1.14196E-14 | 81.28 | P:GO:0006259; F:GO:0003676 |
| KE372074.1_607750 | retrotransposon protein, putative, Ty3-gypsy subclass | 250 | 6.58833E-15 | 84.04 | P:GO:0006259; F:GO:0003676; F:GO:0004518; F:GO:0016740 |
| KE371752.1_119001 | kinesin-like protein KIN-14I | 250 | 1.35358E-15 | 100 | P:GO:0009987; F:GO:0000166; F:GO:0003774; F:GO:0003824; F:GO:0005515; C:GO:0005737;  C:GO:0005856 |
| KE372791.1_92280 | retrotransposon protein, putative, Ty3-gypsy subclass | 250 | 2.60546E-16 | 80.44 | P:GO:0006259; F:GO:0003676;  F:GO:0004518 |
| KE370914.1_71643 | retrotransposon protein, putative, unclassified | 250 | 1.30758E-17 | 84.76 | P:GO:0006259; F:GO:0003676 |
| KE373008.1_174771 | retrotransposon protein, putative, unclassified | 250 | 1.12767E-17 | 92.5 | P:GO:0006259; F:GO:0003676; F:GO:0004518; C:GO:0005575 |
| KE373068.1_467211 | retrotransposon protein, putative, Ty1-copia subclass | 250 | 3.98365E-18 | 81.74 | P:GO:0006259; F:GO:0003676 |
| KE372063.1_625886 | retrotransposon protein, putative, Ty1-copia subclass | 250 | 3.31283E-18 | 82.25 | P:GO:0006259; F:GO:0003676 |
| KE372722.1_281583 | DExH-box ATP-dependent RNA helicase DExH9 | 250 | 5.72496E-19 | 100 | P:GO:0006139; P:GO:0009056; F:GO:0000166; F:GO:0003723;  F:GO:0016787 |
| KE372833.1_339080 | retrotransposon protein, putative, Ty1-copia subclass | 250 | 6.39275E-20 | 84.63 | P:GO:0006259; F:GO:0003676 |
| KE371074.1_138120 | retrotransposon protein, putative, Ty3-gypsy subclass | 250 | 1.04847E-20 | 83.07 | P:GO:0006259; F:GO:0003676;  F:GO:0004518 |
| KE371710.1_188907 | retrotransposon protein, putative, unclassified | 250 | 6.79109E-21 | 83.72 | P:GO:0006259; F:GO:0003676 |
| KE373204.1_177063 | ubiquitin-activating enzyme E1 3 | 250 | 5.93868E-21 | 93.02 | P:GO:0006464; P:GO:0006950; P:GO:0009056; F:GO:0000166; F:GO:0003824; C:GO:0005737 |
| KE372561.1_82783 | retrotransposon protein, putative, unclassified | 250 | 1.68193E-21 | 85.18 | P:GO:0006259; P:GO:0009058; P:GO:0019538; F:GO:0003676; F:GO:0004518; F:GO:0016740 |
| KE372083.1_314445 | retrotransposon protein, putative, Ty1-copia subclass | 250 | 7.93652E-22 | 94.36 | P:GO:0006259; F:GO:0003676 |
| KE372647.1_114388 | retrotransposon protein, putative, Ty1-copia subclass | 250 | 1.2207E-22 | 80.52 | P:GO:0006259; F:GO:0003676 |
| KE372835.1_1132896 | retrotransposon protein, putative, unclassified | 250 | 4.45884E-23 | 88.1 | P:GO:0006259; F:GO:0003676; F:GO:0004518; C:GO:0005575 |
| ASSH01051858.1_1160 | retrotransposon protein, putative, unclassified | 250 | 1.83881E-24 | 82.09 | P:GO:0006259; F:GO:0003676; F:GO:0004518; C:GO:0005575 |
| KE372901.1_192474 | putative polyprotein | 250 | 2.73765E-27 | 86.19 | P:GO:0006259; F:GO:0003676 |
| KE370772.1_207896 | retrotransposon protein, putative, Ty1-copia subclass | 250 | 3.19993E-28 | 85.29 | P:GO:0006259; F:GO:0003676 |
| KE372018.1_1374293 | retrotransposon protein, putative, unclassified | 250 | 1.39409E-28 | 84.92 | P:GO:0006259; P:GO:0009058; F:GO:0003676; F:GO:0004518; F:GO:0016740; C:GO:0005575 |
| KE373023.1_410699 | retrotransposon protein, putative, Ty1-copia subclass | 250 | 5.20178E-29 | 83.05 | P:GO:0006259; F:GO:0003676 |
| KE370807.1_50729 | retrotransposon protein, putative, Ty1-copia subclass | 250 | 1.72699E-29 | 89.27 | P:GO:0006259; F:GO:0003676 |
| KE372735.1_319233 | retrotransposon protein, putative, Ty1-copia subclass | 250 | 1.17353E-29 | 84.49 | P:GO:0006259; F:GO:0003676 |
| KE371507.1_812350 | retrotransposon protein, putative, Ty1-copia subclass | 250 | 6.78892E-30 | 84.91 | P:GO:0006259; F:GO:0003676 |
| KE373101.1_109853 | wall-associated receptor kinase 3 | 250 | 7.7561E-31 | 99.5 | P:GO:0006464; P:GO:0007165;  F:GO:0000166 |
| KE373334.1_302626 | retrotransposon protein, putative, Ty1-copia subclass | 250 | 3.39195E-31 | 84.55 | P:GO:0006259; F:GO:0003676 |
| KE372345.1_416953 | retrotransposon protein, putative, Ty1-copia subclass | 250 | 1.00732E-31 | 89.58 | P:GO:0006259; F:GO:0003676 |
| KE371163.1_615293 | retrotransposon protein, putative, Ty1-copia subclass | 250 | 5.21957E-32 | 93.9 | P:GO:0006259; F:GO:0003676 |
| KE371441.1_774150 | retrotransposon protein, putative, Ty1-copia subclass | 250 | 4.0713E-32 | 94.53 | P:GO:0006259; F:GO:0003676 |
| KE370997.1_839364 | retrotransposon protein, putative, Ty1-copia subclass | 250 | 3.71169E-32 | 89.56 | P:GO:0006259; F:GO:0003676 |
| KE371751.1_53815 | retrotransposon protein, putative, Ty3-gypsy subclass | 250 | 1.04999E-32 | 90.9 | P:GO:0006259; F:GO:0003676; F:GO:0004518; C:GO:0005575 |
| KE373033.1_118064 | retrotransposon protein, putative, Ty1-copia subclass | 250 | 4.79557E-34 | 91.65 | P:GO:0006259; F:GO:0003676 |
| KE372173.1_1448351 | BTB/POZ domain-containing protein At2g13690 | 250 | 8.17395E-35 | 80.79 | P:GO:0006464 |
| KE372760.1_299511 | retrotransposon protein, putative, Ty1-copia subclass | 250 | 1.62008E-35 | 95.52 | P:GO:0006259; F:GO:0003676 |
| KE372978.1_40257 | hypothetical protein DAI22_01g289000 | 250 | 7.08602E-08 | 92.59 |  |
| KE373245.1_1811459 | loricrin | 250 | 1.60946E-09 | 84.49 |  |
| KE373212.1_729142 | BEACH domain-containing protein C2 | 250 | 6.50353E-12 | 83.98 |  |
| KE372660.1_235175 | vWA domain containing protein | 250 | 7.09787E-19 | 83.33 |  |
| KE372871.1_139162 | Stem-specific protein TSJT1, putative, expressed | 250 | 1.82279E-23 | 84.04 |  |
| KE372561.1_48905 | uncharacterized protein LOC112939671 | 250 | 5.64517E-25 | 89.5 |  |

Table S4: comparison of mean environmental conditions between groups of population using two-sample *t*-test allowing for unequal variances

| Variable | Units | N | S | *t*-value | *d.f.* | *P-*value |
| --- | --- | --- | --- | --- | --- | --- |
| Wet-dry frequency | days/year | 7.9 | 13.46 | -8.314 | 8 | 0.001* |
| bio3 | bio2/bio7 | 24.5 | 27.75 | -1.155 | 8 | 0.3122 |
| bio5 | °C × 10 | 295.67 | 323.25 | -2.429 | 4 | 0.0594 |
| bio8 | °C | 233.5 | 258.75 | -1.132 | 8 | 0.294 |
| bio15 | CV | 99.83 | 59.25 | 5.431 | 7 | 0.0029* |
| Soil type |  | 17.83 | 41.75 | -3.563 | 5 | 0.016* |
| UVB2 | J/m 2 /day | 1.34E+05 | 1.24E+05 | 1.552 | 6 | 0.2184 |
| Soil PH | pH | 6.754 | 5.924 | 2.715 | 9 | 0.038* |
| SOC | g/cm 3 | 5.579 | 5.004 | 1.187 | 9 | 0.300 |
| Elevation | m | 69 | 35 | 1.374 | 6 | 0.2067 |

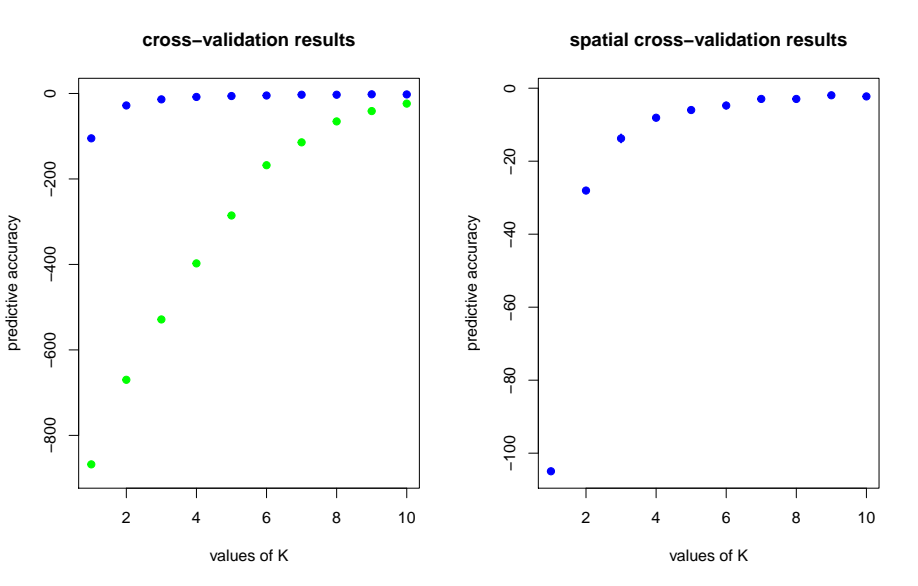
Figure S1: Spatial and non-spatial predictive-accuracy test for the clustering of populations indicating presence of IBD and thus support for spatial model

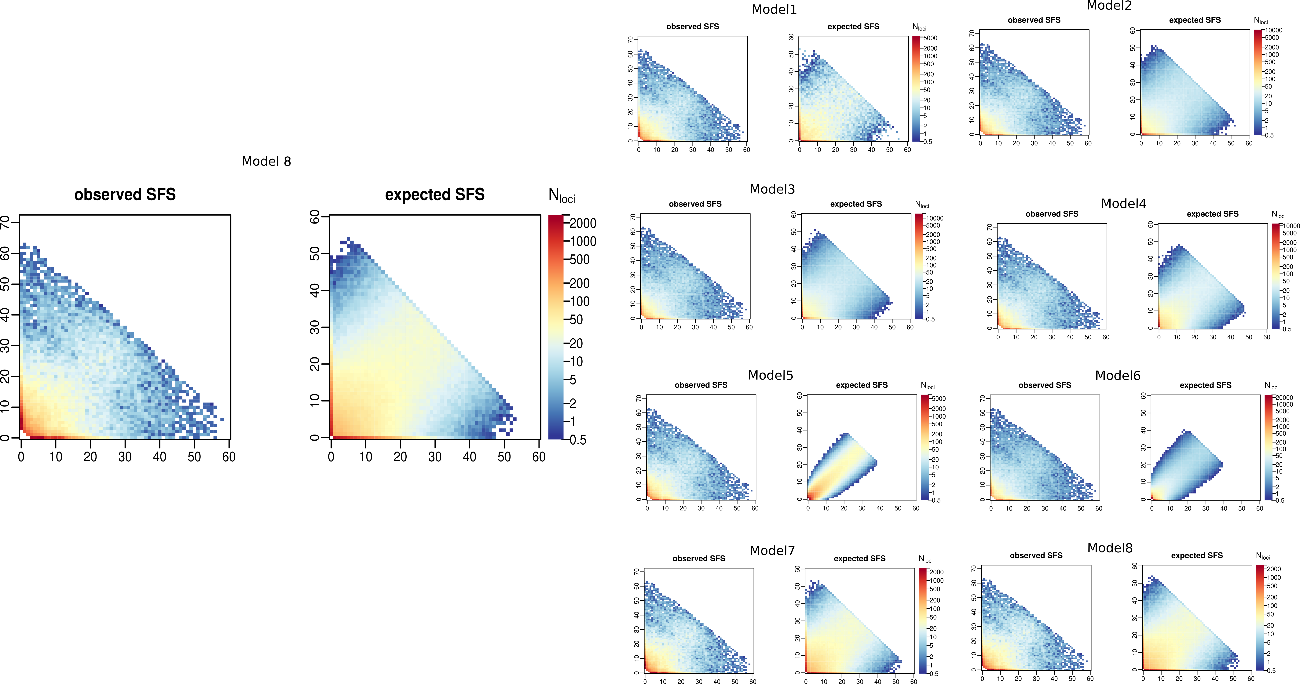

Figure S2: Plots of simulated sfs and sfs from our sequence data indicating that model eight with or without geneflow is the best models and that model 5-6 is poorly supported indicating that there is no possibility that the population size is increasing

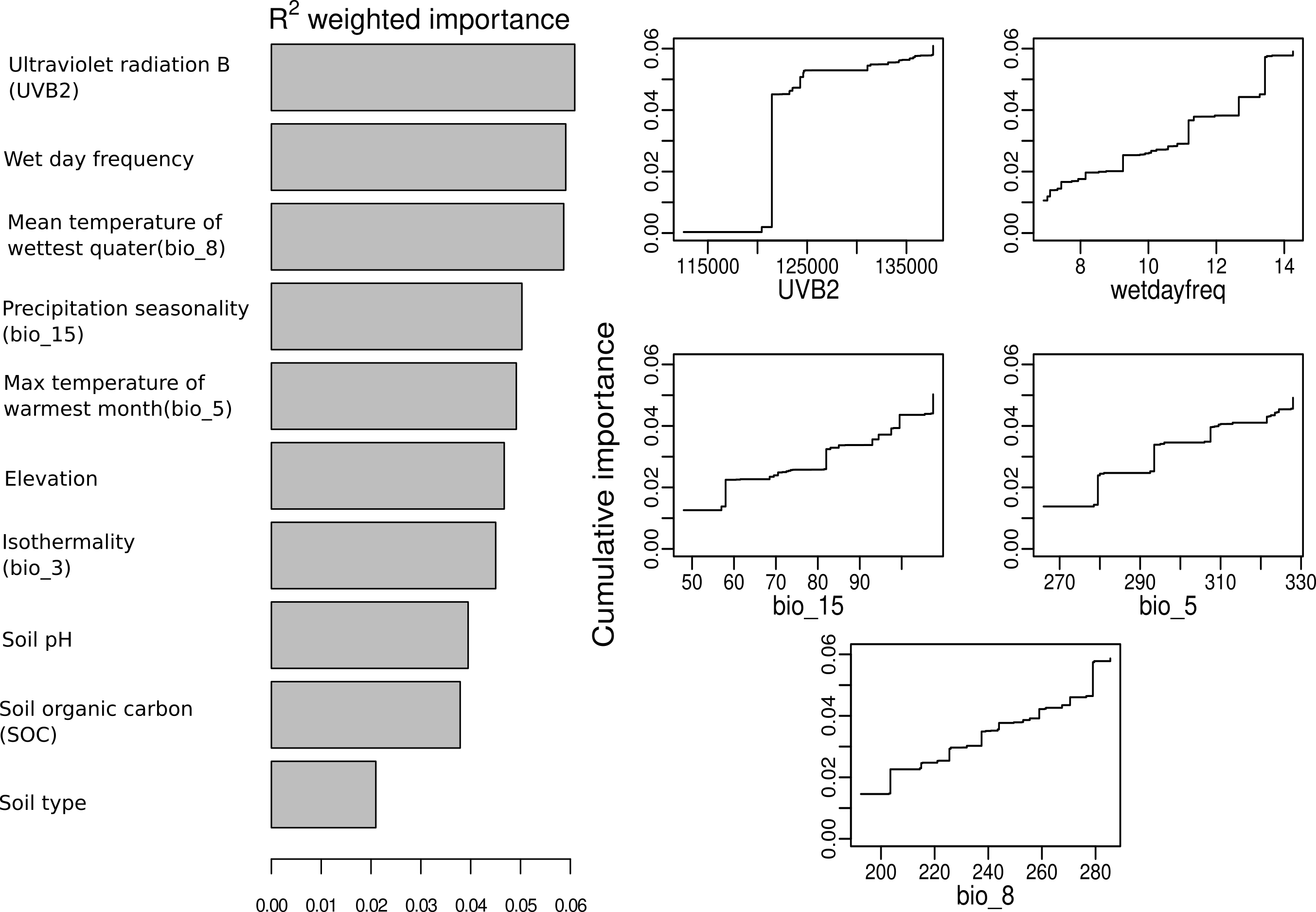

Figure S3: Gradient Forest analysis showing the 5 most important environmental variables explaining variation in allele frequency for the pruned SNPs

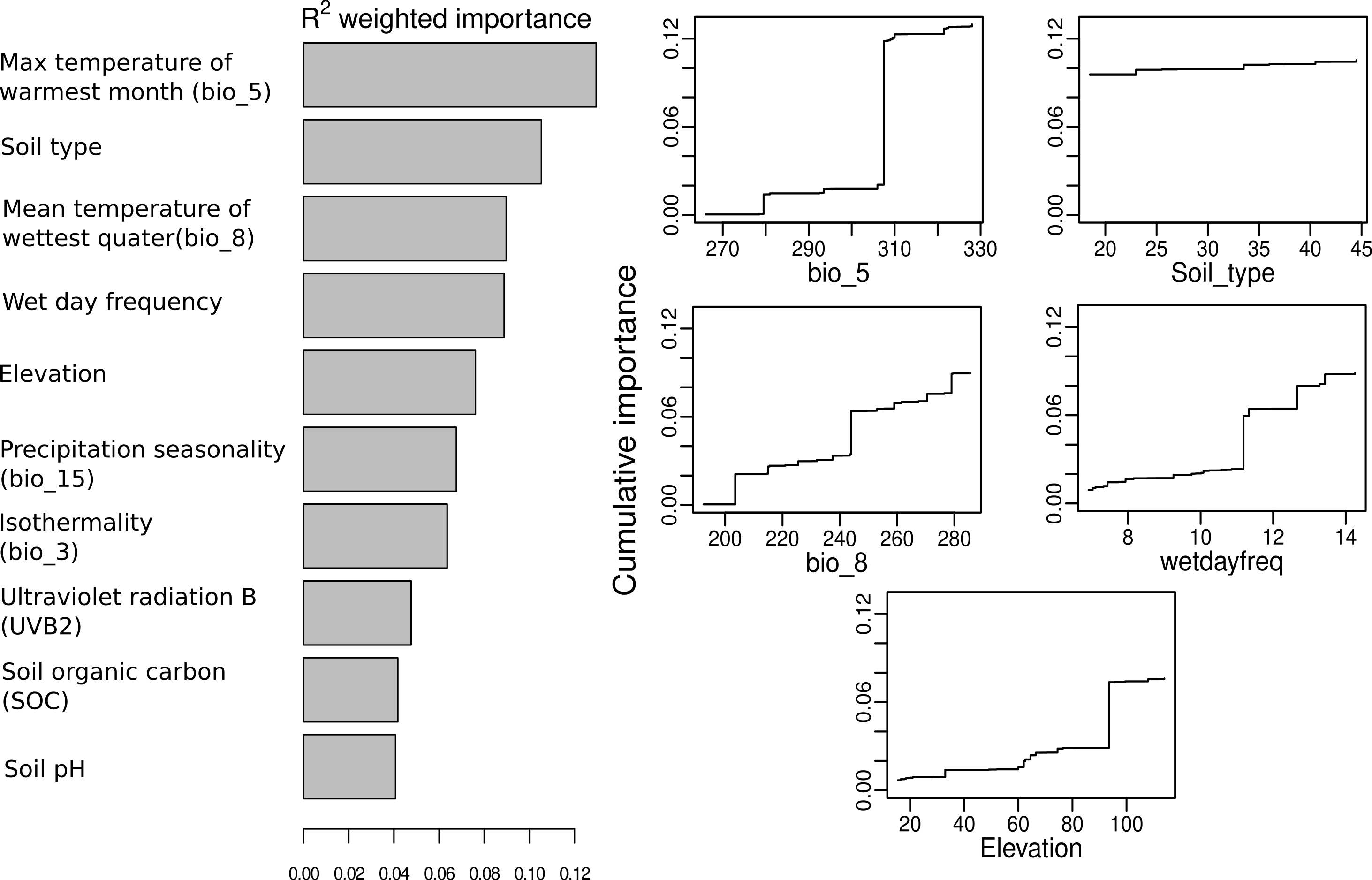

Figure S4: Gradient Forest analysis showing the 5 most important environmental variables explaining variation in morphology

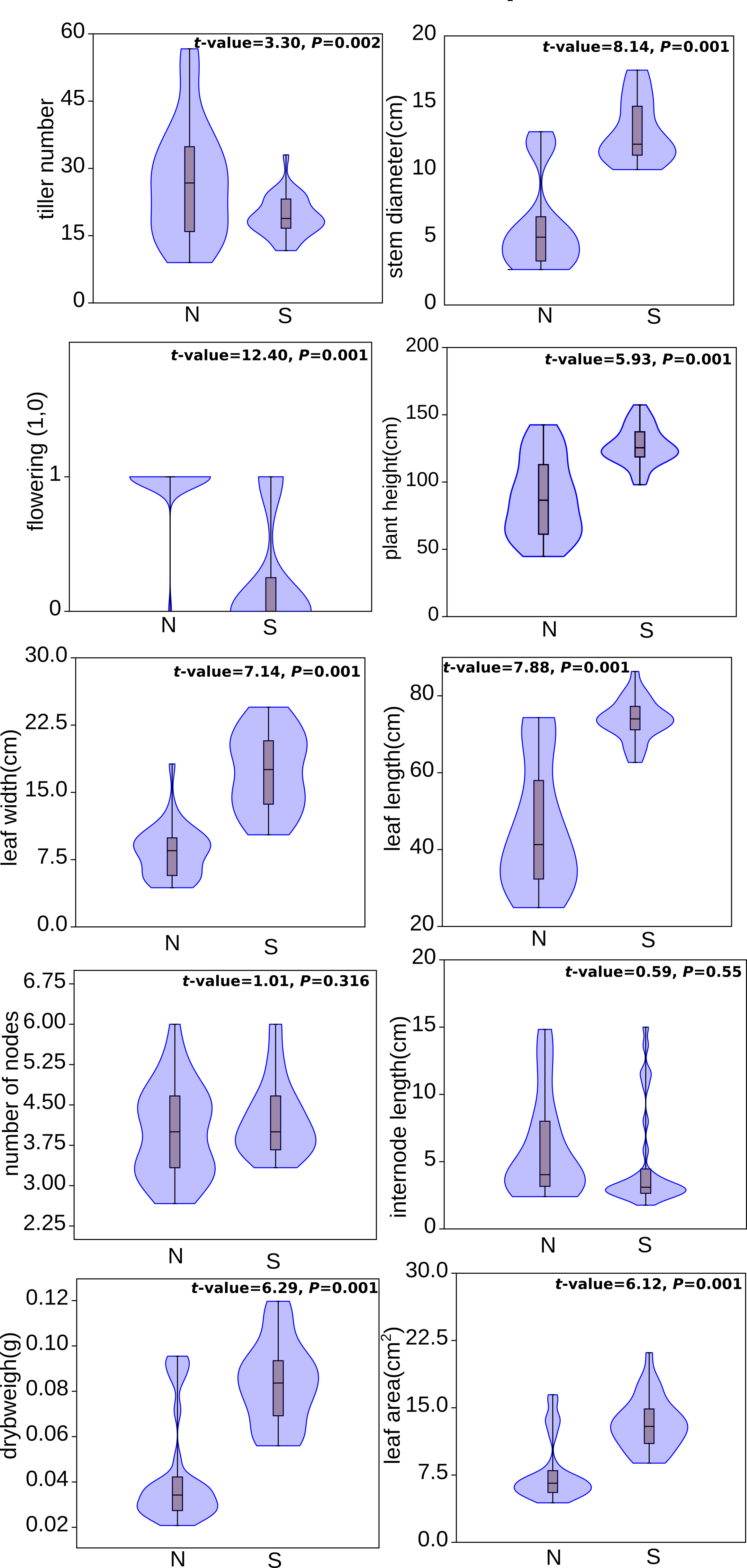

Figure S5: Trait variation between the two genetic groups (North clade, N and South clade, S) using a *t*-test

Figure S6: Phylogenetic diversity using all loci, pruned neutral loci and adaptive loci
